# To the homeRNAmax: Developing an Improved Blood Self-Collection and Stabilization Platform for Remote Transcriptomic Studies

**DOI:** 10.1101/2025.11.24.690053

**Authors:** Madeleine Eakman, Filip Stefanovic, Jean Berthier, Ingrid H. Robertson, Liam A. Knudsen, Serena H. Nguyen, Cosette A. Craig, Kelsey M. Leong, Damon Wing Hey Chan, Karen N. Adams, Ayokunle O. Olanrewaju, Sanitta Thongpang, Tristan M. Nicholson, Erwin Berthier, Ashleigh B. Theberge, Amanda J. Haack

**Affiliations:** Department of Chemistry, University of Washington, Seattle, Washington 98195, United States; Department of Mechanical Engineering, University of Washington, Seattle, Washington 98195, United States; Department of Bioengineering, University of Washington, Seattle, Washington 98195, United States; Department of Urology, School of Medicine, University of Washington, Seattle, Washington 98195, United States; Department of Occupational and Environmental Health Sciences, School of Public Health, University of Washington, Seattle, Washington, 98195, United States; School of Medicine, University of Washington, Seattle, Washington 98195, United States

**Author notes:** Address correspondences to Ashleigh B. Theberge,; Amanda J. Haack. ME and FS contributed equally to this work.

## Abstract

Shifting human subjects research from research sites to participants’ homes removes barriers to participation, including transportation and scheduling difficulties. Previously, we developed homeRNA, a kit for immediate stabilization of RNA in self-collected blood using a custom-engineered tube containing RNA stabilizer fluid. The stabilized RNA is extracted and used for downstream gene expression analysis. Here, we introduce homeRNAmax, which improves our original design by interfacing with a commercially available blood collection tube (BD Microtainer), allowing homeRNAmax to be used with any blood collection method that uses this tube and doubling the possible sample volume that can be collected and stabilized compared to the original homeRNA. Through a pilot study (n=19 participants), we show that homeRNAmax (with the Tasso+ blood collection device) produces RNA samples of sufficient quality (mean RIN=7.8) and yield (mean yield=1.93 μg) for downstream analysis and can reach participants across the United States, who generally (n=17/19) found the homeRNAmax kit easy to use. A key aspect of the homeRNA and homeRNAmax platforms is a fluidic feature that prevents the RNA stabilizer from spilling, however our previous work had not yet fully characterized the mechanism of this feature. Here, we developed a theoretical model of the spill-resistant feature. In brief, fluid in the tube is suspended due to a balance of pressures; an increase in air volume within the tube reduces the air pressure above the fluid, creating a small vacuum, and preventing fluid leakage. Overall, we show that homeRNAmax is a user-friendly, effective tool for remote blood RNA stabilization.

## Introduction

Gene expression analyses using RNA are established methods of monitoring health and disease over time, allowing for immunological studies of infectious diseases and cancers^1–5^. Blood is commonly used for such studies, but blood collection has historically presented challenges for participation in transcriptomics studies because venous blood draws have to be performed by a trained phlebotomist in a clinic, leading to barriers caused by scheduling, aversion to needles, or proximity to a clinic. Shifting sample collection from a clinic to a participant’s home therefore addresses barriers to participation in transcriptomics studies.

Dried blood spots (DBS) are an established method of blood sampling for biomolecule analysis and are an integral tool for remote studies^6–8^. DBS are a very important part of the remote study toolkit^9–13^; however, degradation of RNA after collection– especially in elevated temperature conditions–can limit accurate gene expression profiling^8,14,15^. Hence, we saw the need for a technology that is robust to elevated temperatures and varied shipping conditions.

To this end, we previously developed a custom-designed platform that facilitates the stabilization of RNA in whole blood samples collected at home. The homeRNA kit^16^ makes frequent longitudinal sample collection logistically less challenging compared to clinic-based studies by using a lancet-based upper-arm blood collection device (Tasso, Inc.), removing the need for a trained phlebotomist and the need to milk the collection site (as would be required with DBS sampling using a finger stick). The collected blood is stabilized using a custom-engineered stabilizer tube which holds RNA*later*, a commercially available RNA stabilizing agent. homeRNA has been successfully used to investigate gene expression related to SARS-CoV-2 infection^17,18^ and wildfire exposure^19^, and has been shown to be robust in some high temperature climates and shipping scenarios^20,21^. While our lab uses RNA*later* (ThermoFisher Scientific) to stabilize RNA in blood, homeRNA can be used to stabilize other bioanalytes such as DNA by using a different stabilizer. homeRNA was designed to interface specifically with the Tasso-SST blood tube, as selection of upper-arm blood collection devices was limited at the time of its invention. The recent discontinuation of the Tasso-SST and replacement with the Tasso+ and Tasso Mini presents an opportunity and a need to update homeRNA for broader usability with blood collection tools and downstream analyses.

Here, we introduce the homeRNAmax platform and characterize the spill-resistant feature of the stabilizer tube with a theoretical model. homeRNAmax improves upon the original design by interfacing with the commercially available BD Microtainer blood collection tube instead of a device-specific tube, allowing homeRNAmax to be used with multiple blood collection devices such as the Tasso+ (Tasso, Inc.), the Tasso Mini (Tasso, Inc.), the TAP® Micro Select (YourBio Health), and the RedDrop One (RedDrop Dx). The BD Microtainer also holds up to 1 mL of blood and is available with many coatings, including dipotassium ethylenediaminetetraacetic acid (K_2_EDTA) and lithium heparin, doubling the possible sample volume compared with the original homeRNA and opening up a new range of possibilities for downstream analyses and a range of new study designs to probe questions previously out of reach.

## Experimental details

### Development and assembly of the homeRNA blood collection and stabilization kit

#### Design process for the RNA stabilizer tube

Like the original homeRNA, the RNA stabilizer tube was assembled from three components: 1) a reagent vial to house the liquid stabilizer 2) an adapter that interfaces both the reagent vial and the BD Microtainer (BD) and 3) a vial closure cap. Preliminary design iterations using the same dimensions of the spill-resistant feature as in the original homeRNA^16^, but with an expanded volume to accommodate additional stabilizing fluid, were generated on Solidworks and 3D printed using a Form 3B 3D printer (Formlabs) and a CADworks ProFluidics 285D 3D printer (CADworks). These preliminary designs were evaluated for leakage and spillage using a simulation of the mechanical agitation of the shipping process; three assembled tubes were filled with 3.6 mL RNA*later* (Thermofisher Scientific), then inverted three times for three seconds each. They were then dropped from approximately three feet three times, and were shaken in a shipping box for 30 seconds. Designs were considered to have “passed” this drop test if minimal spillage occurred in observation (maximum of a few droplets of fluid, approximately 20 μL). Specific features of the stabilizer tube design include a cone nozzle feature in the adapter (Fig. 1B) to prevent contact between the stabilizer reagent and the user during blood stabilization (the “spill-resistant” fluidic feature), and an aperture at the top with dimensions to allow the BD Microtainer to fit snugly with the homeRNAmax setup without making the removal of the cap difficult for the user. Engineering drawings of the stabilizer tube components can be found in the Supplementary Information (Fig. S1).

**Figure 1.**
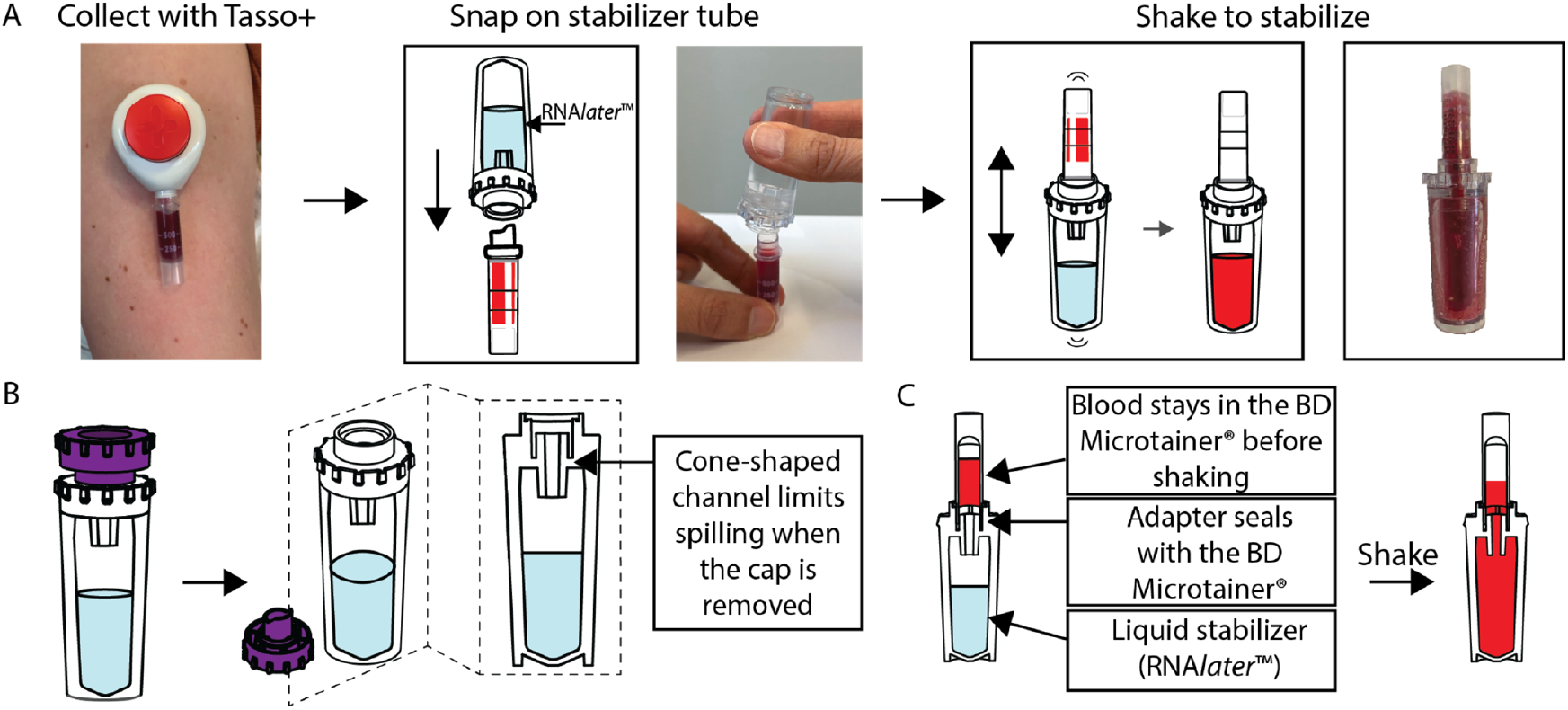
Workflow and features of homeRNAmax. A) describes the workflow for RNA stabilization of blood samples using homeRNAmax. To use our system, participants collect blood by following the directions for the Tasso+ device, connecting the BD Microtainer blood tube to the stabilizer tube filled with RNA*later*, then shake the blood into the RNA*later* to stabilize the RNA in the blood sample. B) Schematic of filled stabilizer tube. The features shown here are adapted from our original homeRNA system reported in Haack, Lim et al.^16^, with adjustments made to interface with BD Microtainers. C) Cross section schematic of the stabilizer tube interfacing with a blood-filled BD Microtainer before and after shaking to stabilize.

### Design considerations for injection molding of the RNA stabilizer tube

The adapter and the vial components were injection molded out of polycarbonate (PC: Makrolon 2407), and the vial cap was injection molded out of polypropylene (PP: RTP Permastat 100 Anti-static) by Protolabs, Inc. (Maple Plain, MN). Several design considerations were optimized to include features important for the process of injection molding. Briefly, these include 1) Adding at least a 0.5 degree draft to all vertical surfaces in the adapter and the vial to aid in removal from the mold, (the part of the cap that interfaces with the adapter was left undrafted to ensure a tight seal with the adapter), 2) flipping the direction of draft partway down the adapter to allow easier removal of both sides of the mold, 3) thickening of the vial and adapter pieces to allow for ejector pin zones, 4) removing sections of plastic in the cap to prevent warping, and 5) designing the adapter piece to avoid any overhangs, allowing for a two-piece mold. The vial and the adapter pieces were both made of polycarbonate to ensure consistent material shrinkage, and because polycarbonate is a common material for biochemical samples containers. The cap was made out of polypropylene because the added flexibility of the polypropylene mimics the BD Microtainer and helped ensure a tight seal between the cap and the adapter.

### Fabrication of the RNA stabilizer tube

For assembly, all components of the stabilizer tube were injection molded by Protolabs, Inc. as described above and cleaned via sonication in 70% ethanol (v/v) for 30 minutes and air dried. The adapter was bonded onto the reagent vial using a UV curing glue (Dymax MD® 1450-M-UR-SC) (Fig. S2). Bonded parts were cured for 45 minutes at 405 nm in a Formcure (Formlabs). The stabilizer tube was filled with 3.6 mL of RNA*later* as the stabilizing reagent, capped, and checked for leakage due to bonding defects before distribution to study participants.

### Measuring evaporative loss from stabilizer tubes

The stabilizer tube was assembled according to the section above (see *Fabrication of the RNA stabilizer tube*) before being filled with 3.6 mL of RNA*later*. Since the manufacturer recommended ratio is 1 mL of blood to 2.6 mL of RNA*later*, 3.6 mL of RNA*later* was chosen to accommodate any loss due to evaporation or shipping, as well as collected blood volumes above 1 mL, ensuring stabilization in a variety of scenarios. The tubes were sealed with the injection molded caps and their initial mass was recorded. The filled stabilizer tubes were then placed upright and kept on a lab benchtop. The mass of the stabilizer tube was then recorded once per week for a total of 10 weeks (Fig. S3). The change in mass from initial mass is calculated and converted to volume, using a density of 1.2125 g/mL for RNA*later*. The resulting volume is then subtracted from 3.6 mL and converted to a percentage, representing the percent RNA*later* loss from evaporation for each tube at each timepoint.

### Development of the stabilizer tube theoretical model

The theoretical model was prepared using MATLAB and Microsoft Excel.

For testing the stabilizer tubes for leakage, four stabilizer tubes were filled with 2.00 mL, 4.00 mL, 6.00 mL, and ∼6.70 mL (completely filled) with DI water. The height of the fluid within the tube was measured using a caliper, and all the tube nozzles were dried using a Kimwipe. All tubes were then inverted, and the heights of the fluid above the nozzles were measured using a caliper. Tubes were observed for leakage for 10 minutes.

### User experience pilot study

This study was IRB approved under STUDY00007868. For the pilot study, we recruited participants using Facebook ads. We selected individuals who had not previously seen or used a Tasso device (by self-report) and who were not affiliated in any way with the Bioanalytical Chemistry for Medicine and the Environment (BCME) lab (Theberge lab). Individuals were eligible if they were healthy, adults aged 18 or older, and able to have lancet blood sampling (collected by self-report). Prospective participants were screened for age, gender, race and ethnicity, and state of residence. Balancing these factors to ensure our study population is representative of that of the United States, people who were selected were sent an email with a link to the informed consent form (ICF). Upon completion of the ICF, participants would be considered as “enrolled” and were instructed to complete a survey with their shipping information. The study kit consisted of (1) instructions for sampling and sample return (Fig. S4-S5), (2) a Tasso+, (3) Tasso warming pad, (4) an alcohol wipe, (5) a bandaid, and (6) a homeRNAmax stabilizer tube which was filled with 3.6 mL of RNA*later* on the day of shipment (Fig. S6 and Table S1). The kits were packed inside a United Parcel Service (UPS) LabPak alongside another, pre-labeled UPS LabPak for sample return. The packages were sent using the UPS next day air service and the tracking information was directly shared with the participants. Once a package was delivered to a study participant, another email with study instructions (see SI) was sent that included a link to an instructional video prepared by our lab. The participants then completed their sampling one time according to the instructions and completed a short usability survey (See SI). The sample was then sealed back in the Tasso box, placed into the spare LabPak, and kept indoors overnight. The following morning, participants left their samples at the designated pickup location and the study team scheduled a UPS pickup. Once the samples arrived back to the lab, they were stored at −20°C until further processing.

### Blood collection and RNA stabilization

In the pilot study outlined above, participants self-collected blood using the Tasso+ device, which is commercially available and collects up to 1 mL of blood. The Tasso+ device uses BD Microtainers for blood collection. In the pilot study, we opted to use the K_2_EDTA coated BD Microtainer tube to limit blood coagulation prior to stabilization. Participants were instructed to collect blood for five minutes or when the blood reached the collar of the BD Microtainer (1 mL), whichever happened first; see instructions for use (IFU) in the Supplemental Information (Fig. S4) for more information. Once the blood was collected, participants connect the BD Microtainer to the stabilizer tube containing 3.6 mL of RNA*later* and shake them to mix the blood and stabilizer.

### RNA isolation and quantification

Sample RNA extraction was performed using the blood RiboPure RNA Purification Kit and following the manufacturer suggested protocol. Because homeRNAmax sample volumes exceeded the maximum capacity for this kit, the samples were split into two aliquots and extracted at the same time. Prior to the elution step, these two aliquots were combined into a single sample to have a uniform sample for analysis. The resulting RNA was then analyzed using the Agilent 2100 Bioanalyzer (Agilent Technologies, Inc.) to verify RNA integrity. The RNA yield was measured both on the BioTek Cytation 5 (Agilent Technologies, Inc.) Take3 application, as well as on the Qubit Flex Fluorometer (Thermo Fisher Scientific Inc.) for comparison.

## Results

The original homeRNA^16^ was designed to accommodate the Tasso-SST blood tube, which could only hold up to 500 μL of blood. Our new design interfaces with the commercially available BD Microtainer tube, which holds up to ∼1 mL of blood. By interfacing with the BD Microtainer, homeRNAmax can be used with the Tasso+, Tasso Mini, and other blood collection devices that use the BD Microtainer as its blood tube, such as the YourBio TAP® Micro Select.

Several design considerations were made to ensure homeRNAmax performs comparably to the original homeRNA^16–18,20,21^. The dimensions of the cuvette had to be adjusted to hold at least 5 mL of fluid to accommodate the new blood volume of 1 mL, the RNA*later* required to stabilize 1 mL of blood, and extra RNA*later* to safeguard against issues such as evaporation or participants collecting additional drops of blood such that the total blood volume is >1 mL (see Experimental Details for more information).

The homeRNAmax cuvette can reasonably hold up to ∼6 mL of total volume (up to ∼5 mL of stabilizer with ∼1 mL of sample) as a consideration for applications other than RNA stabilization of blood that might require a larger volume. The homeRNAmax dimensions are also designed such that stabilized samples (i.e., the cuvette and adapter assembly connected to the blood tube) fit in a 50 mL conical vial, providing an easily accessible commercially available method to give additional protection for lab technicians and shipping personnel (although strictly speaking this is beyond the secondary containment required for shipping blood samples, which include only a watertight secondary container with absorbent material and a rigid box) The cuvette and adapter assembly is also short enough for current commercially available plastic pasteur pipettes to reach the bottom, allowing for the entire sample volume to be extracted without the risks associated with using glass pasteur pipettes.

Considerations were also taken to allow for injection molding, including drafting vertical surfaces, avoiding overhang, and thickening the vial and adapter components to allow for ejector pin placement. These considerations are described in detail in the Experimental Details section.

### Theoretical model of stabilizer tube

To understand the mechanism underlying the spill-resistant nozzle design on the homeRNAmax stabilizer tube (Table S2), we developed a model of pressures acting upon the liquid-air interface in the middle or bottom of the nozzle. Initially, fluid is added to the stabilizer tube when it is in an upright position (Fig. 2A). Upon inversion, the liquid is suspended in the middle (Fig. 2Bi) or bottom (Fig. 2Bii) of the nozzle when there is a balance of pressures at the liquid-air interface (i.e., *P*_*balance*_=0). Inside the tube, we consider the hydrostatic pressure (*P*_*hyd*_), the pressure of the trapped air (*P*_*air*_), a capillary pressure at the interface between the liquid and the atmosphere within the nozzle (*P*_*cap*_), and—if the liquid is at equilibrium at the bottom of the nozzle—a pressure of the fluid pinning to the opening of the nozzle (*P*_*pin*_). The capillary pressure of the upper surface of the liquid is negligible since the tube diameter is larger than the capillary length. On the outside of the tube, the only pressure is the atmospheric pressure (*P*_*atm*_). The balance of pressures can be expressed using the following equations in the case where the meniscus is suspended within the nozzle (1) and the case where the meniscus is pinned at the nozzle opening (2), respectively:

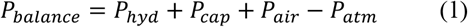

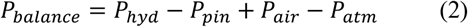

**Figure 2.**
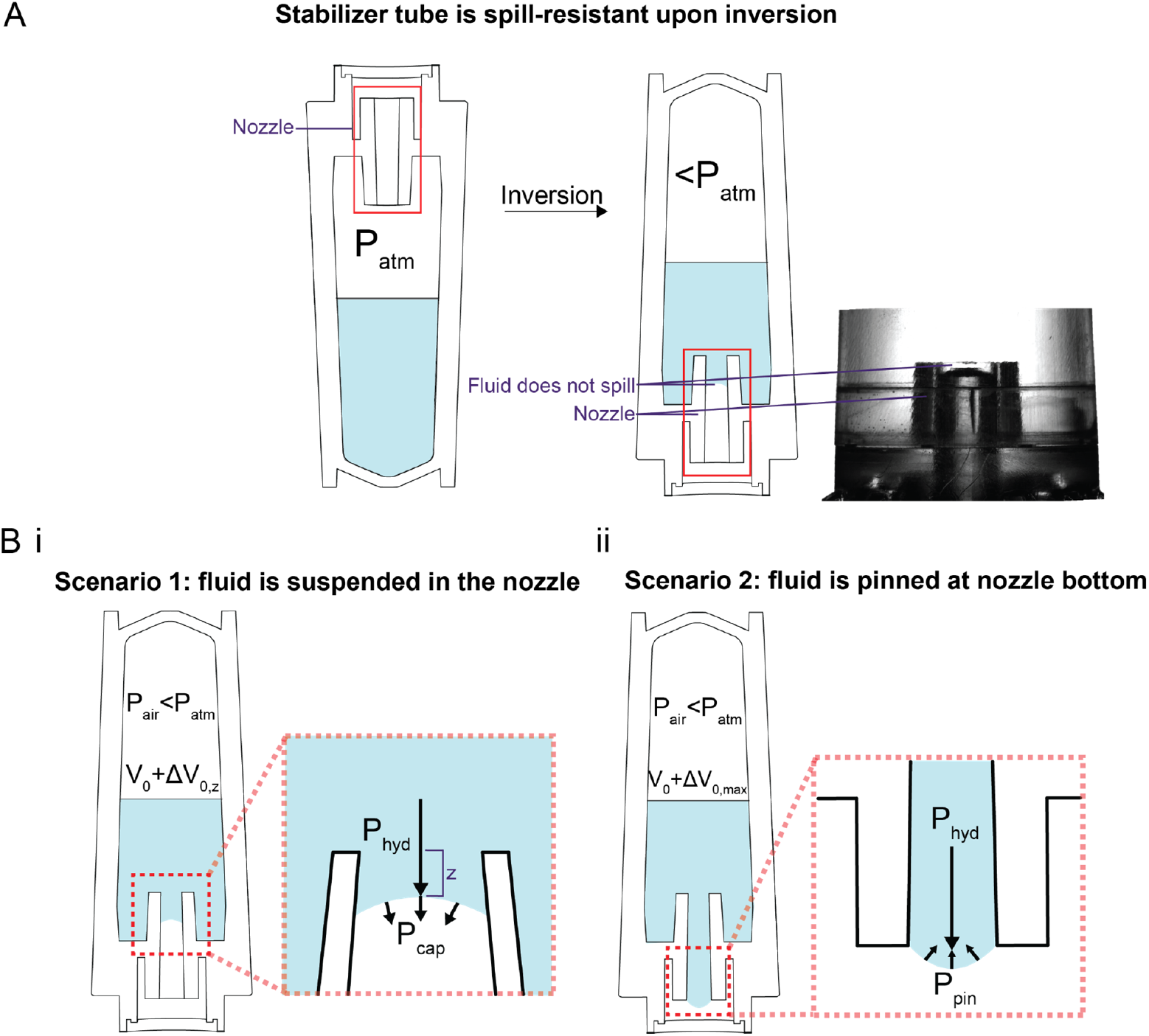
homeRNAmax stabilizer tube is spill-resistant due to a balance of pressures within the nozzle. (A) Schematic of stabilizer tube before (left) and after inverting (right). A close-up image of an inverted homeRNAmax tube showing a suspended meniscus within the nozzle is depicted to the right of the schematic. (B) Schematic of an inverted stabilizer tube at equilibrium when i) fluid stops in the nozzle and ii) fluid pins to the nozzle opening at equilibrium. The former is observed for the dimensions and volumes used in this work.

The notation in equations (1) and (2) is that the values for the individual pressures are always positive; the signs of each pressure indicate whether their contributions are upwards or downwards. The most important pressure for preventing spilling is the trapped air pressure, which decreases as the tube is inverted and is, as a result, lower than the atmospheric pressure. The expansion of the trapped air occurs when a small amount of fluid moves into the nozzle, lowering the pressure inside the tube.

This drop in pressure can be expressed using an equation derived from the Ideal Gas Law (3):

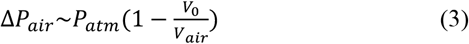

where *V*_*O*_ is the initial volume of air (i.e., the air volume before inversion), and *V*_*air*_ is the volume of the air after tilting. In the scenario described in Fig. 2Bi, upon inverting the tube, the pressure decreases with the increasing air volume associated with the fluid entering the nozzle (equivalent to the volume of fluid in the nozzle, *ΔV*_0,*z*_), and creates a vacuum strong enough that the fluid is suspended in the middle of the nozzle. In this case, the drop in pressure opposes the hydrostatic and capillary pressures, arresting the meniscus within the nozzle and preventing spillage. This is the scenario that was observed experimentally for the dimensions of the stabilizer tube depicted in this work.

The most unfavorable situation for preventing leakage is when the meniscus is located at the bottom of the nozzle (Fig. 2Bii) after tilting the tube. This scenario was never experimentally observed with the homeRNAmax device in the dimensions and volumes used in this work. However, it was observed using 3D printed stabilizer tubes where the height was increased to ∼17 cm (as opposed to 4.8 cm for our actual stabilizer tube used in homeRNAmax) and the draft was removed. In these 3D printed tubes and at low liquid volumes, the effects of fluid movement into the nozzle resulted in less meaningful pressure changes. In this situation depicted in Fig. 2Bii, pinning holds the fluid at the bottom of the nozzle, and the fluid leaks when pinning is exceeded (when the pendant drop exceeds a hemisphere). For further details on the development of the theoretical model, see Supplemental Information Appendix 1.

In some cases over timescales on the order of 10-20 minutes, additional meniscus movement has been observed after reaching equilibrium; we postulate that this movement is a result of effects related to temperature (e.g., evaporation, temperature changes over time). We have not observed any leakage as a result of these effects. Furthermore, in actual use cases, the tube is not left uncapped and inverted for longer than times on the order of seconds.

### User Experience Pilot Study

To validate the usability and robustness of the homeRNAmax platform, we conducted a human subjects pilot study of a total of 20 participants. For the pilot study, we were able to recruit a diverse cohort. A total of 319 individuals completed our screening survey, of whom 38 were invited based on the inclusion criteria. From there, 23 signed the informed consent. The discussion below focuses on the 20 participants who completed the sampling and the survey. These participants were located across 20 states (including Hawaii) (Fig. 3A), with a racial background similar to that of the larger United States population (Fig. 3B), and had a fairly even distribution of age and sex (Fig. 3C).

**Figure 3.**
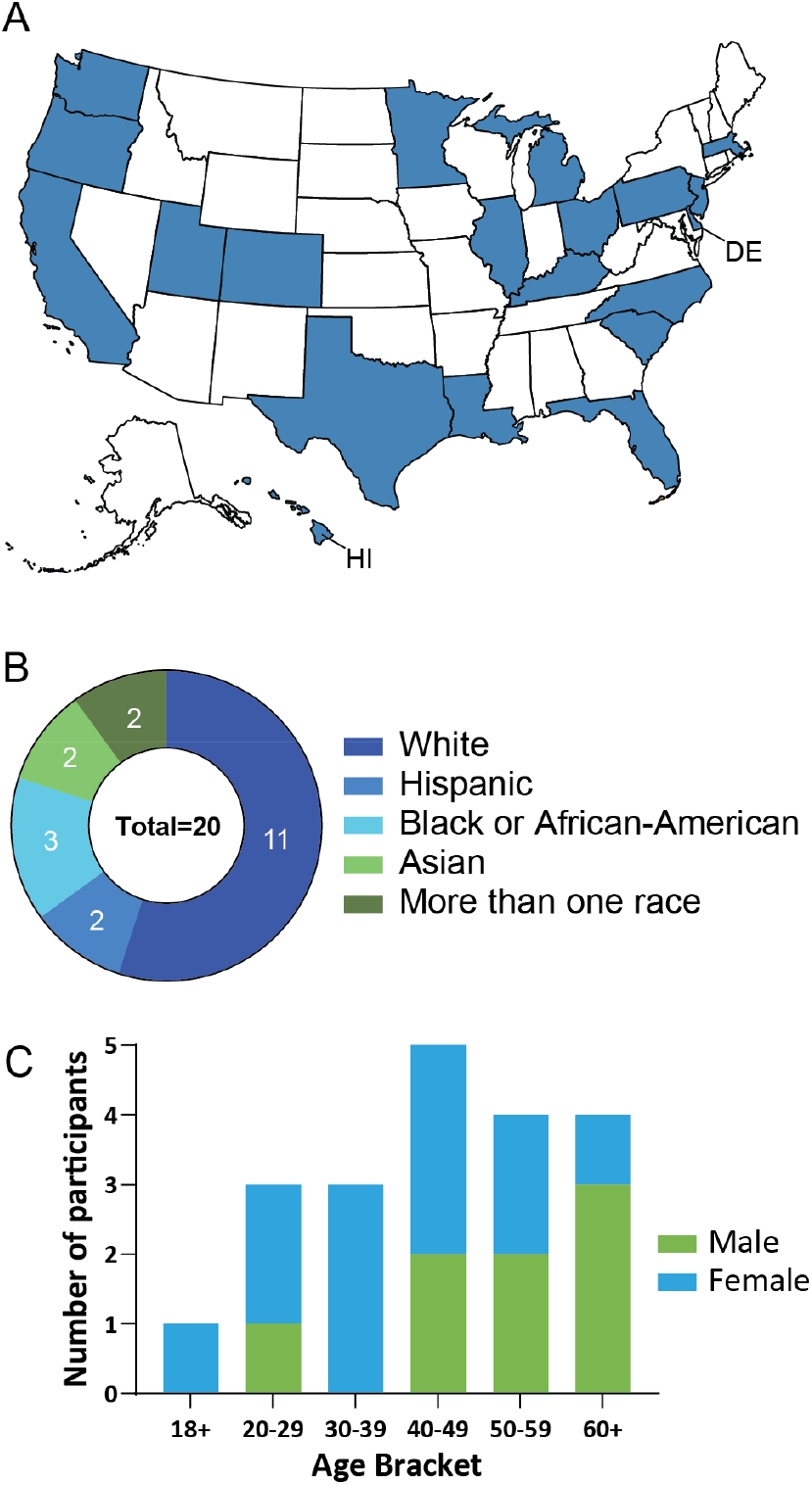
Summary of participant demographics. (A) US map showing distribution of participants across 20 states including Hawaii. Each shaded state represents one participant. (B) Distribution of races and ethnicities of participants. (C) Participant sex and age bracket. Participants were selected to represent an evenly distributed cohort, one that matches as closely as possible to the larger population of the United States.

Each of the 20 participants self-collected and stabilized one blood sample using the homeRNAmax kit. With the sampling, participants were asked to complete surveys about their experience. The majority of the participants (16) indicated that they felt no pain when using the Tasso+ device, and only 4 indicated that they experienced mild pain. (Fig. 4A). No participant reported experiencing moderate or major pain. Additionally, the majority of participants found the Tasso+ device very easy to use (Fig. 4B).

**Figure 4.**
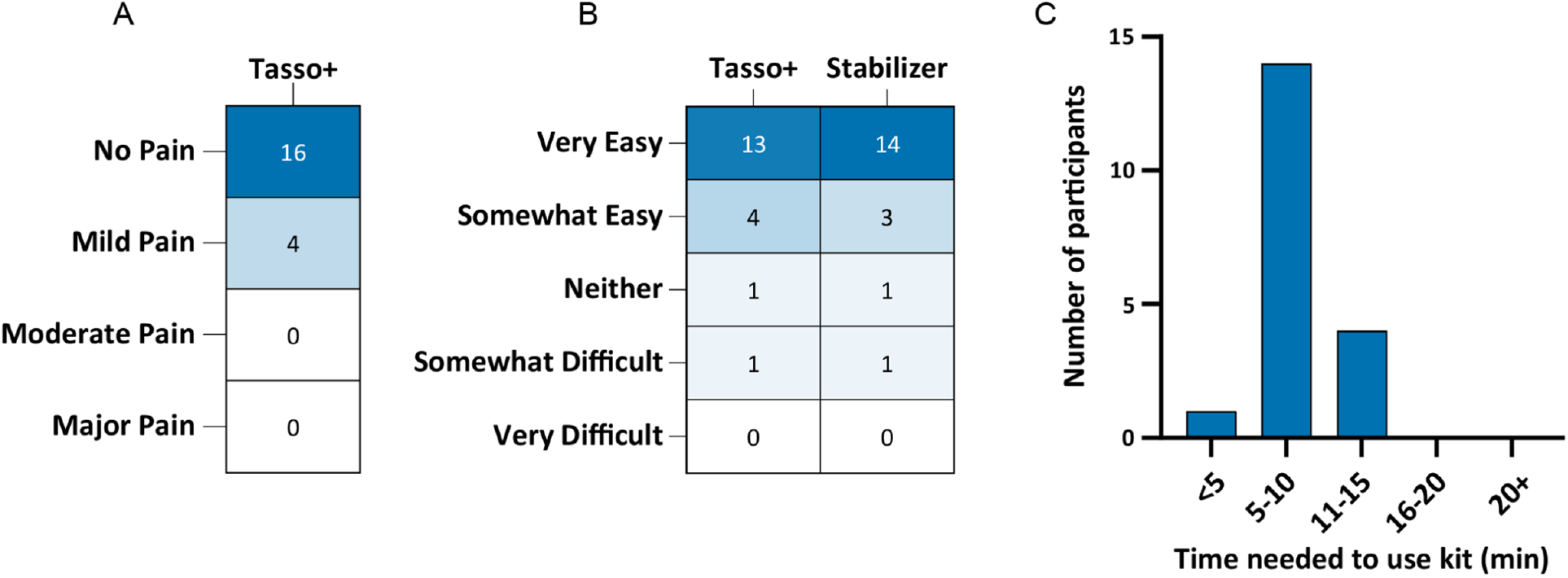
Participant-reported homeRNAmax usability data. (A) Pain level reported by each participant following the use of the Tasso+ device. (B) Participant-reported usability of the Tasso+ and homeRNAmax stabilizer tube following blood stabilization. (C) Time needed to use the kit, including reading instructions, collecting and stabilizing the sample, and packaging sample for pickup.

Of the 20 participants who were enrolled, only one was unable to collect blood. As a result, we did not collect data on usability of the stabilizer tube and blood level of this participant. The remaining 19 participants indicated very similar ease of use of the stabilizer tube (Fig. 4B). Moreover, most of the participants were also able to use the kit within 5 to 10 minutes, making the sampling very time efficient.

Most (n=15/20) participants collected at least 500 μL of blood, and 13 participants collected more than 500 μL. This means that 13 participants (of the 19 who successfully collected blood using the homeRNA platform) were able to collect more blood than the maximum volume of the original homeRNA platform^16^ (Table S2). The homeRNAmax platform, with the Tasso+ device, reliably collected greater volumes of blood than the original homeRNA which used the Tasso-SST, although the majority of participants in our original homeRNA study collected the maximum possible blood volume (Fig. S7).

As with the original device^16,20,21^ validation, we used the Agilent Bioanalyzer 2100 to measure the quality of RNA using the RNA integrity number (RIN). RINs range from 1 to 10, where a RIN of 1 represents a completely degraded sample and a RIN of 10 a perfectly preserved RNA sample^22,23^ (Fig. S8). For this study, we used the Qubit Flex, which is specific to RNA, to measure the total yield of RNA in each of the samples.

Stabilization of homeRNAmax was similar to that of the original homeRNA device, with most (n=16/19) RIN values falling above 7 (Fig. 5A) and most (17 out of 19) samples having a yield of more than 500 ng (Fig. 5B), both of which are common requirements for Illumina RNA sequencing^24–26^. Although the RIN of some of the samples was below 7, which we have also seen in our prior work using the original homeRNA kit^16–21,27^, advances in sample processing, sequencing technologies, and analysis methods can generate readouts for samples with lower RINs and yields^24,28–30^. Bioanalyzer electropherograms and a table of RNA yields can be found in the Supplementary Information (Fig. S8, S9 and Table S3)

**Figure 5.**
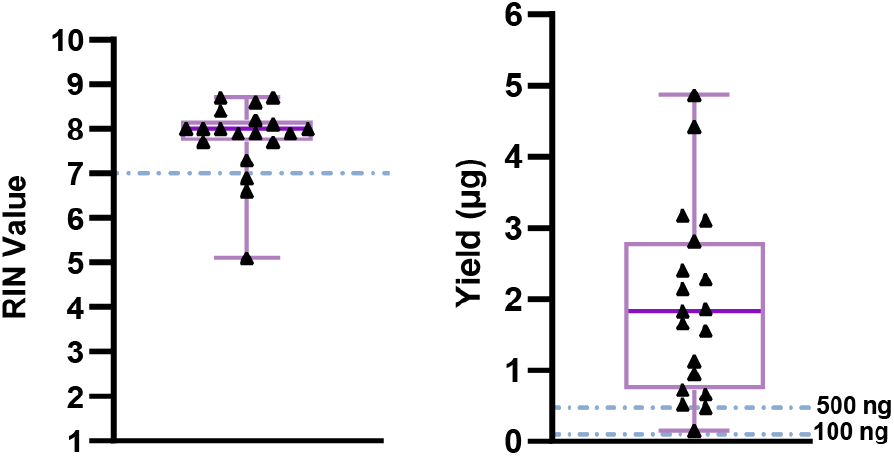
RNA integrity and yield of homeRNAmax-stabilized blood samples from a nationwide remote study. Both graphs are represented as standard box and whisker plots with the triangles representing each sample. (A) RNA integrity number (RIN) of homeRNAmax samples obtained using the Agilent 2100 Bioanalyzer. The dashed line represents a RIN of 7, a conservative cutoff for common RNA sequencing technologies. (B) Yield measurements obtained using Qubit Flex. The dashed line represents 500 ng, a conservative cutoff for yield required for sequencing.

Although homeRNAmax collected more blood than the original homeRNA device^16,20^, the distributions of RNA yields reported are relatively similar. This is expected from our past experiences using homeRNA, where we have seen that blood volume is not necessarily indicative of resulting RNA yield due to large inter-individual variation in RNA per unit volume of blood (for more information, see Fig. S14 of Haack, Lim et al)^16^. However, within a person increased blood volume will still inherently increase RNA collected. When considering other homeRNA studies^16,21^, the median RNA yield tends to fall around 2 and 3 μg (Fig. S10), and the homeRNAmax-stabilized samples fall within that range. It is important to note that the yields reported in the original homeRNA paper were measured with NanoDrop, which uses absorbance of the sample at 260 nm to estimate concentration, and assumes that the sample is pure RNA (has no contamination). From our experience, even with the RiboPure blood RNA extraction kits, there is oftenDNA contamination. The Qubit Flex, which was used to determine the yields in Fig. 5B, relies on a dye that selectively binds to RNA and measures the fluorescence of the bounded dye. As a result, this measurement tool is considered to be more representative of the actual RNA yield in extracted samples. Although the resulting yields from the Take3, which also relies on absorbance, and the Qubit Flex in this study are similar, we note that the Take3 values are slightly higher, as would be expected from a less selective measurement. For comparison of the homeRNAmax platform with the original homeRNA platform, we consider the RNA yield readouts from the Take3 as well as the Qubit Flex. When considering the yields obtained from the Take3 (as a similar measurement method to that used in our original homeRNA manuscript^16^, homeRNAmax performed similarly to or better than the original homeRNA (Fig. S10), further supporting that the RNA yield is not directly tied to collected blood volume. Regardless of the instrument used to measure the yield, nearly all samples had yields higher than the minimum requirement of 500 ng needed for most sequencing technologies.

Additionally, while we have not seen a strong correlation between collected blood volume and resulting RNA yield in our other homeRNA studies (500+ samples^16–21,27^), the increased volume is useful for the RNA extraction process. Because homeRNAmax can have up to twice the sample volume of the original homeRNA, homeRNAmax samples can be aliquotted into two tubes. This allows one aliquot to be kept as whole blood in RNA*later* and banked in the freezer, while the other is extracted. The second banked aliquot can prove to be invaluable in the process of analyzing the data following a human subjects study, providing insurance should the other aliquot be lost due to laboratory error. Additionally, a banked second aliquot can be very useful in longer-term longitudinal studies, where samples are collected on the scale of months to years. In cases like these, it may be desirable to extract some of the earlier time points all together, and then have a second aliquot of the earlier time points that could be extracted later with additional time points. Here, having samples split into two aliquots allows them to be directly compared against another set of samples collected at later time points, while preserving extraction batches for direct comparison. Extracting samples at the same time allows for greater ease of comparison by limiting the differences due to variability between extraction batches. There are many other instances where banking the aliquot may be of use, especially in longitudinal studies, that are dependent on the study design and question being asked.

## Conclusion

homeRNAmax is the next generation of the homeRNA at-home blood self-collection and stabilization kit. It is able to collect up to two times the blood volume of the original kit, making for easier sample handling and enabling banking of RNA-stabilized samples. Like the original homeRNA, homeRNAmax is spill-resistant due to the cone-shaped nozzle geometry, which balances the total pressure of the liquid in the tube to the outside atmospheric pressure upon inversion. In the human subjects pilot feasibility study with a total of 20 participants, homeRNAmax was virtually painless (*n* = 16/20), easy to use (*n* = 13 for Tasso+ blood collection device and *n* = 14 for RNA*later*-containing stabilizer tube), and quick (*n* = 14 only needed 5 - 10 minutes to use the kit). The resulting RNA was shown to be of high quality (mean RIN = 7.8) with most samples above a RIN of 7 and yields above 500 ng, benchmarks often used for most common sequencing technologies.

## Author Contributions

M.E. and F.S. authors contributed equally. All authors have given approval to the final version of the manuscript. *Co-corresponding authors.

## Conflicts of Interest

EB, AJH, and ABT filed patent 17/361,322 (Publication Number: US20210402406A1) and EB, AJH, FS, ME, and ABT filed patent 63/571,012 through the University of Washington on homeRNA and a related technology. ABT reports filing multiple patents through the University of Washington and receiving a gift to support research outside the submitted work from Ionis Pharmaceuticals. EB and ST have ownership in Salus Discovery, LLC, and Tasso, Inc. that develops blood collection systems used in this publication, and is employed by Tasso, Inc. Technologies from Salus Discovery, LLC are not included in this publication. TMN has financial interests/equity in Tasso, Inc. and spouse is employed by Tasso, Inc. EB is an inventor on multiple patents filed by Tasso, Inc., the University of Washington, and the University of Wisconsin-Madison. EB, ST, and ABT have ownership in Seabright, LLC, which will advance new tools for diagnostics and clinical research, including the homeRNA platform used in this publication, and EB and ST are partially employed by Seabright, LLC. The terms of this arrangement have been reviewed and approved by the University of Washington in accordance with its policies governing outside work and financial conflicts of interest in research. AJH, LK, SN, CC, KML, AOO, TMN, and ST have also filed additional patents through the University of Washington.

## Acknowledgements

This publication was supported by the David and Lucile Packard Foundation (for the human subjects component of the research), and the National Institutes of Health (NIH) through the National Institute of General Medical Sciences award number R35GM128648 (for in-lab developments and theory) and R01AR084274 (for device manufacturing). This Research was supported by the National Institute of Environmental Health Sciences K12ES033584 (TMN, trainee).

The REDCap used for human subjects consent and enrollment is supported by the Institute of Translational Health Sciences, which is funded by the National Center for Advancing Translational Sciences of the National Institutes of Health under award number UL1TR002319. The content is solely the responsibility of the authors and does not necessarily represent the official views of the National Institutes of Health or other funding bodies. We would also like to thank the participants.

