## Supplementary Information 1 for "To the homeRNAmax: Developing an Improved Blood Self-Collection and Stabilization Platform for Remote Transcriptomic Studies"

**Table of Contents:**

| | |
|---|---|
| **Figure S1:** Engineering drawings | S2-S4 |
| **Figure S2:** Fabrication of the stabilizer tube | S5 |
| **Figure S3:** Summary of evaporation test data | S5 |
| **Figure S4:** Instructions for Use | S6-S7 |
| **Figure S5:** Stickers illustrating adequate stabilization | S8 |
| **Figure S6:** Components of the homeRNAmax kit | S9 |
| **Table S1:** Components of the homeRNAmax kit | S9 |
| **Table S2:** Summary of data from leakage test | S10 |
| **Table S3:** Summary of blood collection and RNA yield data | S11-S12 |
| **Figure S7:** Comparison of homeRNAmax and original homeRNA blood level | S13 |
| **Figure S8:** General information on Bioanalyzer profile | S14 |
| **Figure S9:** Agilent 2100 Bioanalyzer electropherograms | S15 |
| **Figure S10:** Comparison of RNA yield from present study and original homeRNA study | S16 |
| **Supplementary Text** | S17-S23 |
| **References** | S24 |

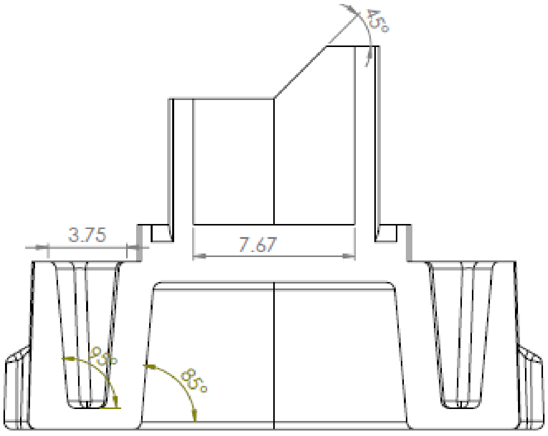


SECTION C-C
SCALE 6 : 1

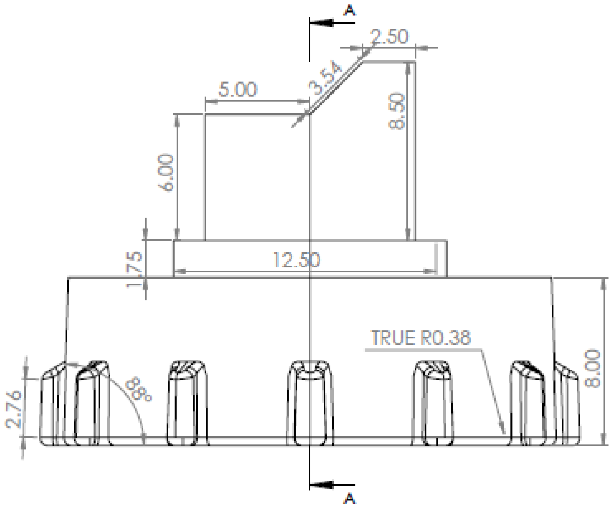


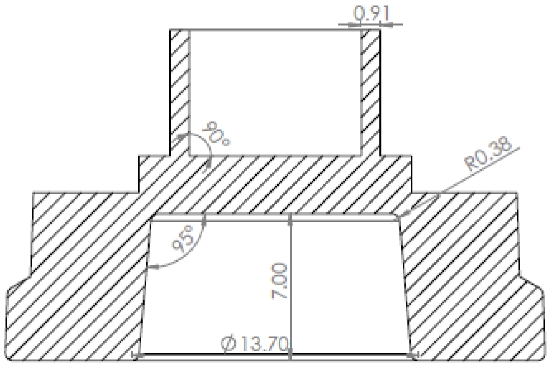


SECTION A-A
SCALE 6 : 1

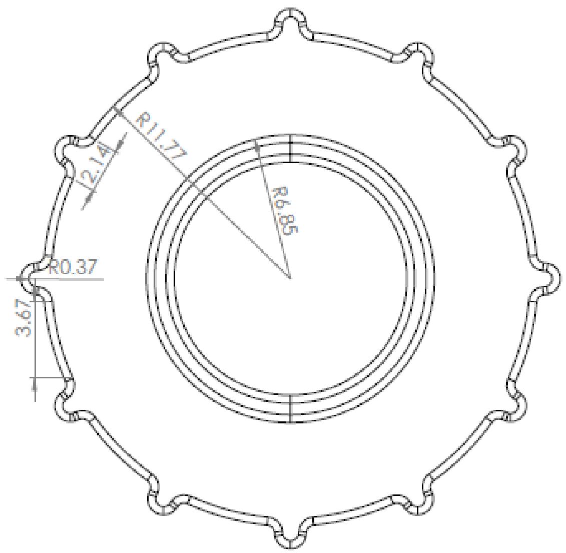


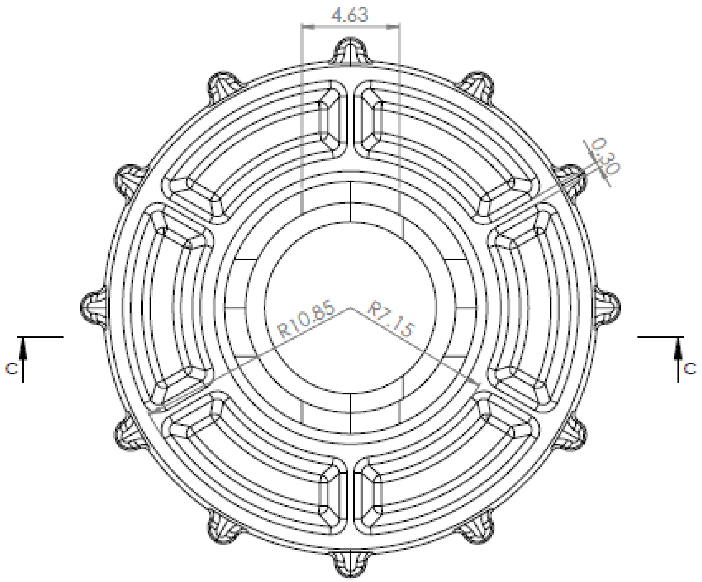

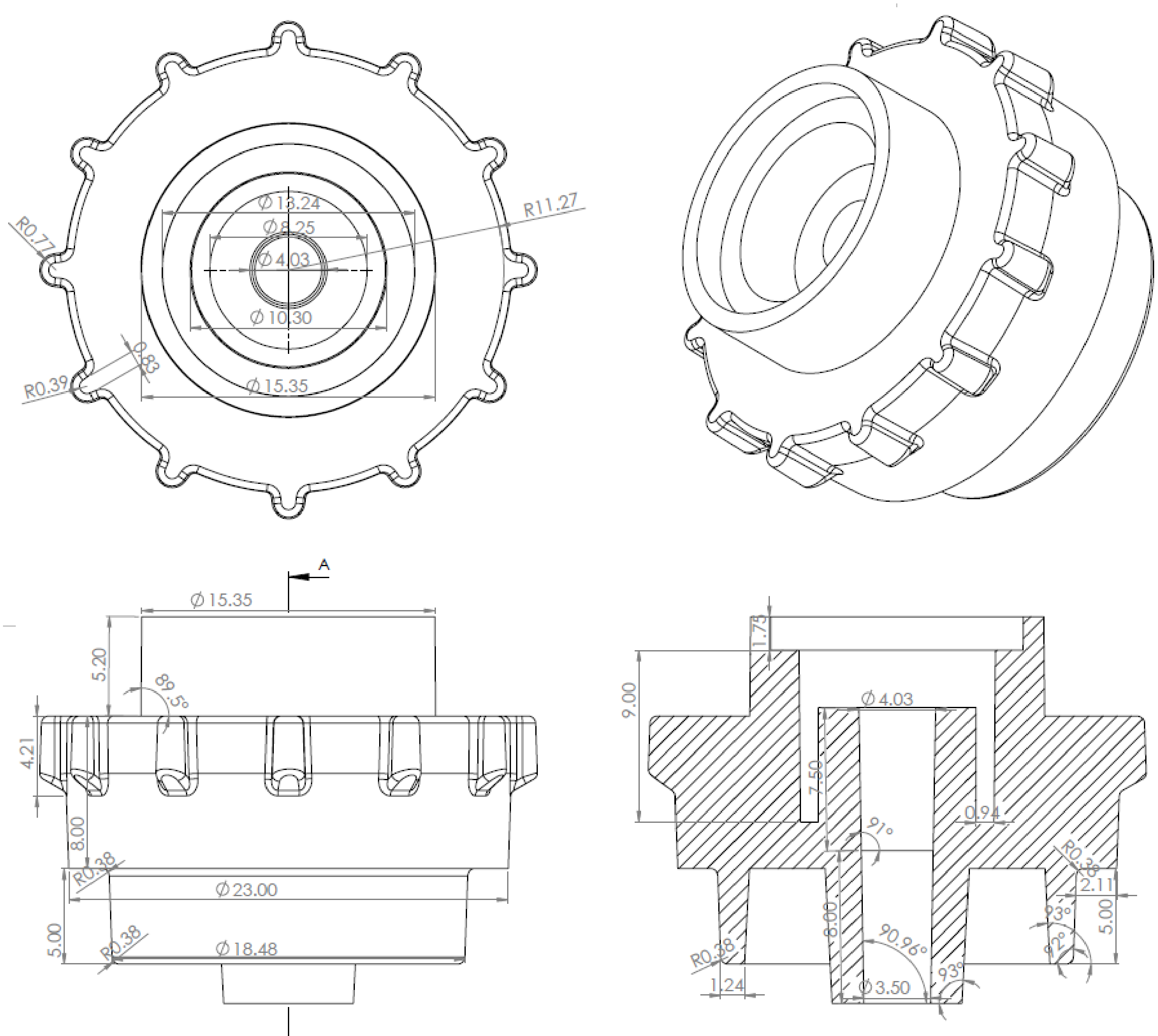


SECTION A-A
SCALE 6 : 1

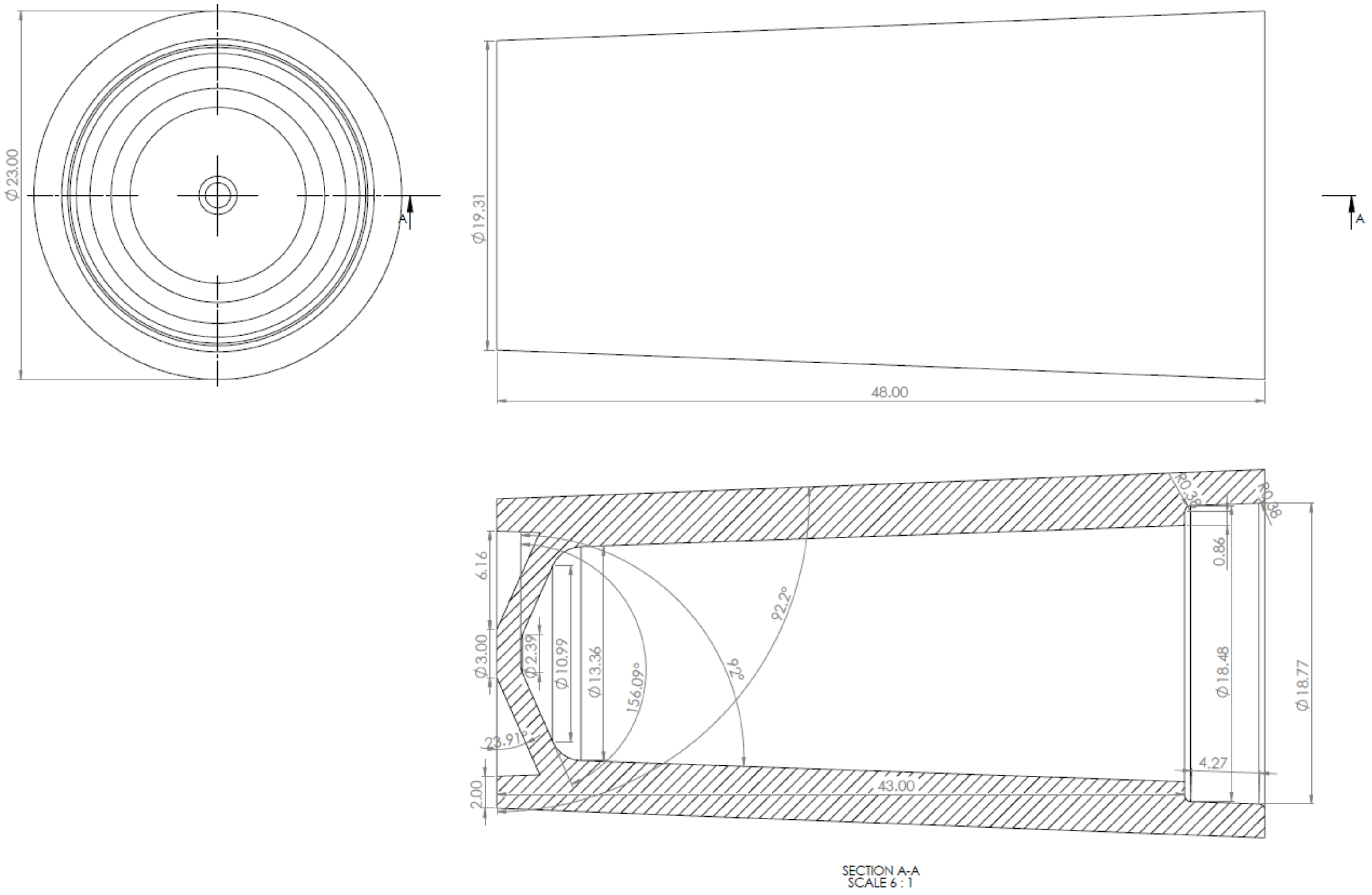


**Figure S1.** Engineering drawings of the homeRNAmax components. All dimensions are in millimeters.

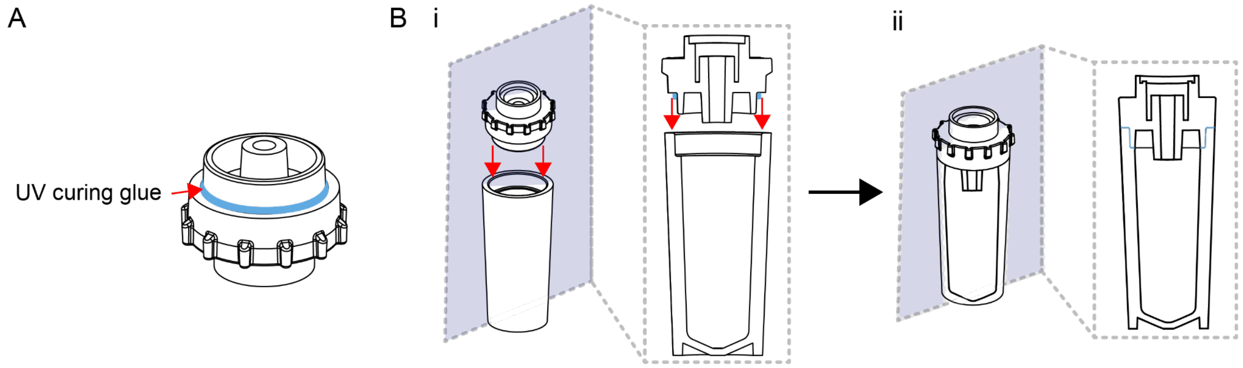


**Figure S2.** Fabrication of stabilizer tube. A) Location where UV curing glue (blue) is applied to the adapter piece. B) Mating of the adapter piece with the vial piece showing location of the glue before (i) and after (ii) inserting the adapter piece into the vial piece. Glue is then cured with a 405 nm UV lamp as described in the materials and methods.

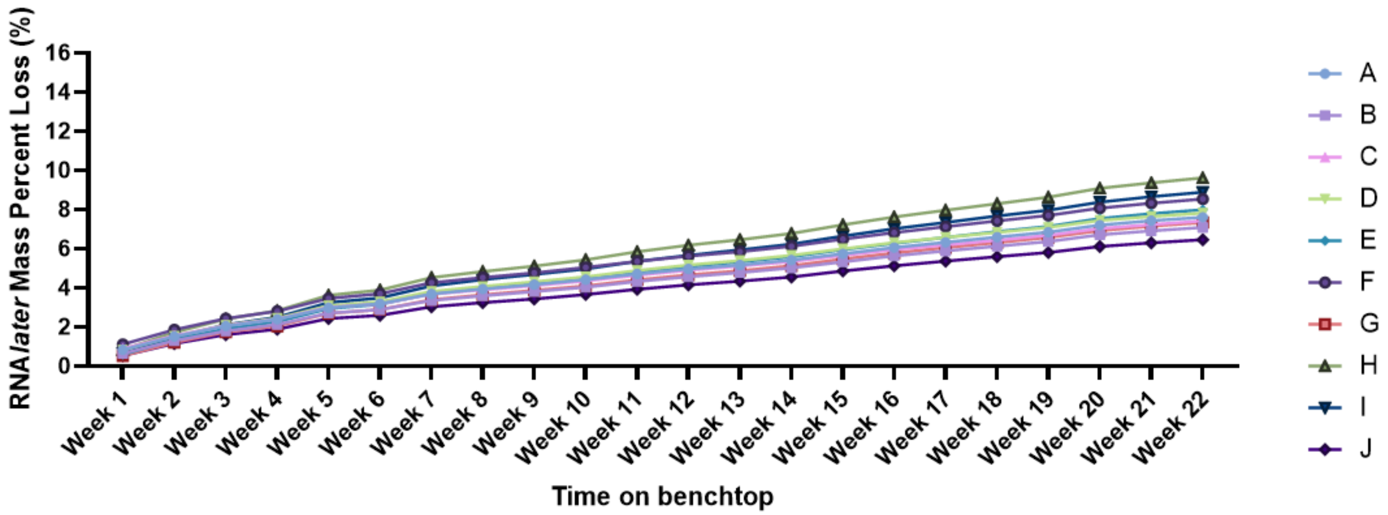


**Figure S3**. Summary of evaporation test data. homeRNAmax tubes were filled with RNA*later* and stored upright at room temperature on a lab bench. The mass was measured each week and a percent loss was calculated. These data show that evaporation is roughly linear with an approximate loss of between 0.5% and 1% (~0.020 - 0.039 grams) per week stored at room temperature.

#### PREPARE FOR COLLECTION

**BEFORE YOU BEGIN**

Point your phone's camera here to watch an instructional video:

Or type this URL:
https://tinyurl.com/ms5eu5f2

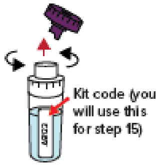


**1. Remove cap from stabilizer tube by twisting it off and discard the cap. Write down the kit code.** Set the tube aside. You will use this in step 12.

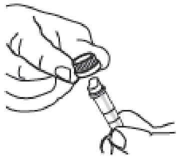

**2. Twist and remove cap from blood collection tube, and discard the cap.**

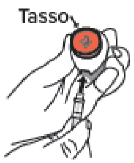


**3. Remove Tasso from box. Connect blood collection tube to Tasso device, and press until snug.**

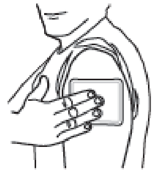

**4. Activate warmer by bending the silver disc. Knead to fully activate, and apply to upper arm for 2 minutes.**

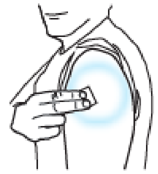

**5. Clean area with alcohol wipe and allow to dry.**

#### COLLECT BLOOD USING TASSO DEVICE

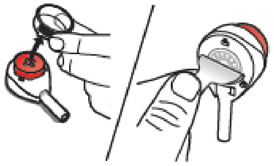

**6. Remove clear plastic cover over the red button on the Tasso device.**

**Peel tab behind the device.**

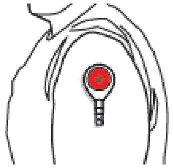

**7. Stick device to upper arm with tube pointing down. Hang arm straight down at side.** Do not remove once it is on.

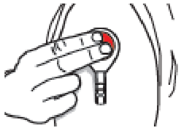

**8. Press button all the way down ONCE and release.** You may not feel anything.

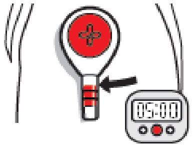

**9. Start a 5 minute timer.** Keep arm at your side. You may not see blood right away. It can take 1-2 minutes for blood to flow. Use a mirror or phone camera as needed to watch blood flow.

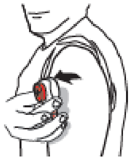

**10. After 5 minutes OR when the blood reaches the collar of the tube, *whichever comes first*, remove the device from your arm.** Many people do not fill the tube to the collar.

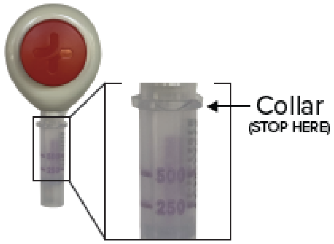


**NOTE: If you have not collected any blood skip to step 15.**

#### STABILIZE AND PACKAGE

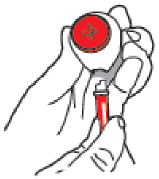

**11. Remove tube by twisting slightly and pulling down.** This may take a bit of finger strength.

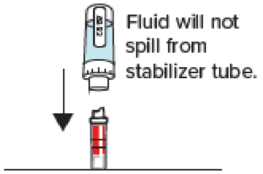


**12. *With the blood collection tube on the bottom*, connect the stabilizer tube (from step 1) and blood tube together until you can't press any farther.**

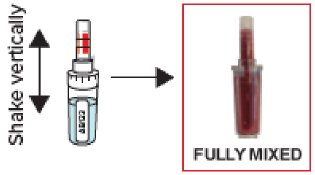


**13. *With the stabilizer tube on the bottom*, shake up and down until the fluid in the top and bottom compartments is the same color. For most participants, this takes about 10 seconds.** If fluid is only in the bottom, it should be the same color throughout.
**DO NOT DISCONNECT TUBES.**

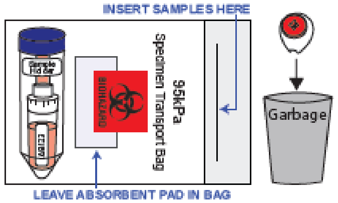


**14. Place blood sample in sample holder, and put into the specimen bag. Place specimen bag inside the original box.** Remaining used materials, including the Tasso device and warmer, can be thrown away.

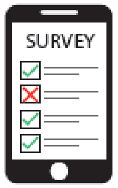


**15. Refer to email instructions for filling out the Online Survey and for mailing your sample back to the lab.**

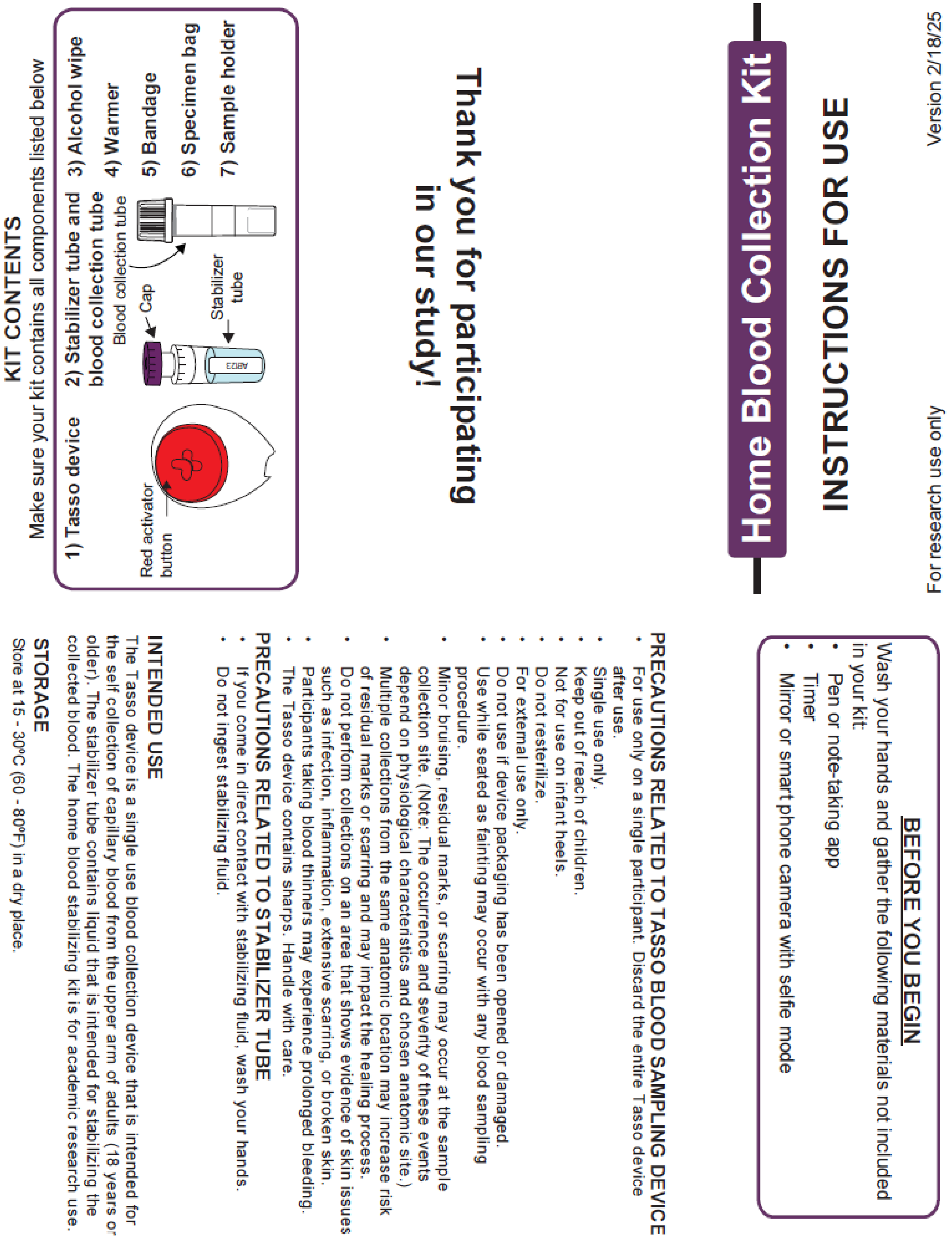

For research use only

Version 2/18/25

### Home Blood Collection Kit

#### INSTRUCTIONS FOR USE

**Thank you for participating in our study!**

##### KIT CONTENTS

Make sure your kit contains all components listed below

1) Tasso device
2) Stabilizer tube and blood collection tube
3) Alcohol wipe
4) Warmer
5) Bandage
6) Specimen bag
7) Sample holder



##### BEFORE YOU BEGIN

Wash your hands and gather the following materials not included in your kit:

- Pen or note-taking app
- Timer
- Mirror or smart phone camera with selfie mode

##### PRECAUTIONS RELATED TO TASSO BLOOD SAMPLING DEVICE

- For use only on a single participant. Discard the entire Tasso device after use.
- Single use only.
- Keep out of reach of children.
- Not for use on infant heels.
- Do not resterilize.
- For external use only.
- Do not use if device packaging has been opened or damaged.
- Use while seated as fainting may occur with any blood sampling procedure.
- Minor bruising, residual marks, or scarring may occur at the sample collection site. (Note: The occurrence and severity of these events depend on physiological characteristics and chosen anatomic site.)
- Multiple collections from the same anatomic location may increase risk of residual marks or scarring and may impact the healing process.
- Do not perform collections on an area that shows evidence of skin issues such as infection, inflammation, extensive scarring, or broken skin.
- Participants taking blood thinners may experience prolonged bleeding.
- The Tasso device contains sharps. Handle with care.

##### PRECAUTIONS RELATED TO STABILIZER TUBE

- If you come in direct contact with stabilizing fluid, wash your hands.
- Do not ingest stabilizing fluid.

##### INTENDED USE

The Tasso device is a single use blood collection device that is intended for the self collection of capillary blood from the upper arm of adults (18 years or older). The stabilizer tube contains liquid that is intended for stabilizing the collected blood. The home blood stabilizing kit is for academic research use.

##### STORAGE

Store at 15 - 30°C (60 - 80°F) in a dry place.

**Figure S4.** Instructions for use for the homeRNAmax kit. Instructional video file and Adobe editable .pdf are included as attachments in the Supplemental Information.

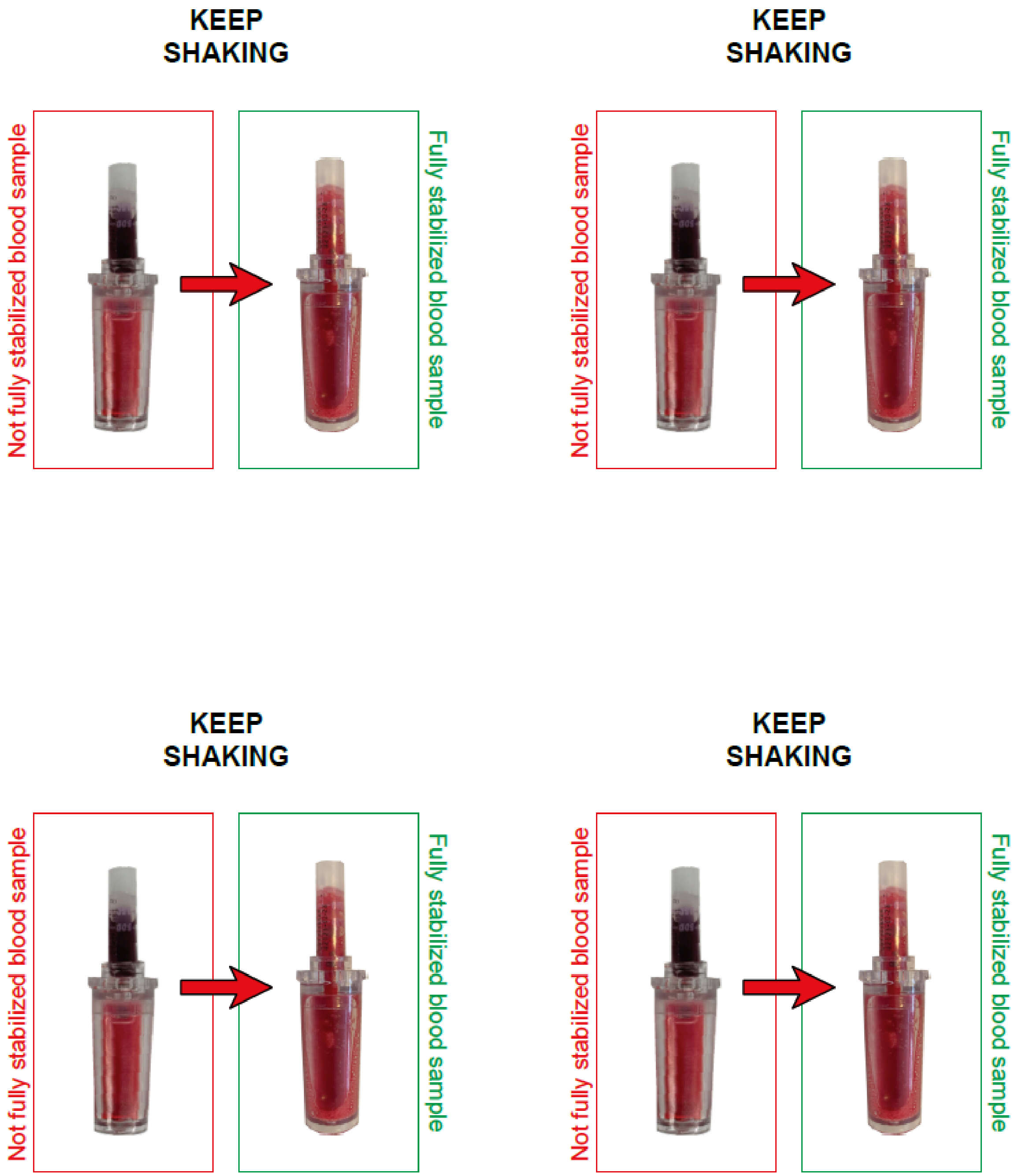


**Figure S5.** Stickers showing adequate stabilization. Stickers adhere to inside of kit box lid. Labels used are Avery 5168. Adobe editable .pdf is included as an attachment in the Supplementary Information.

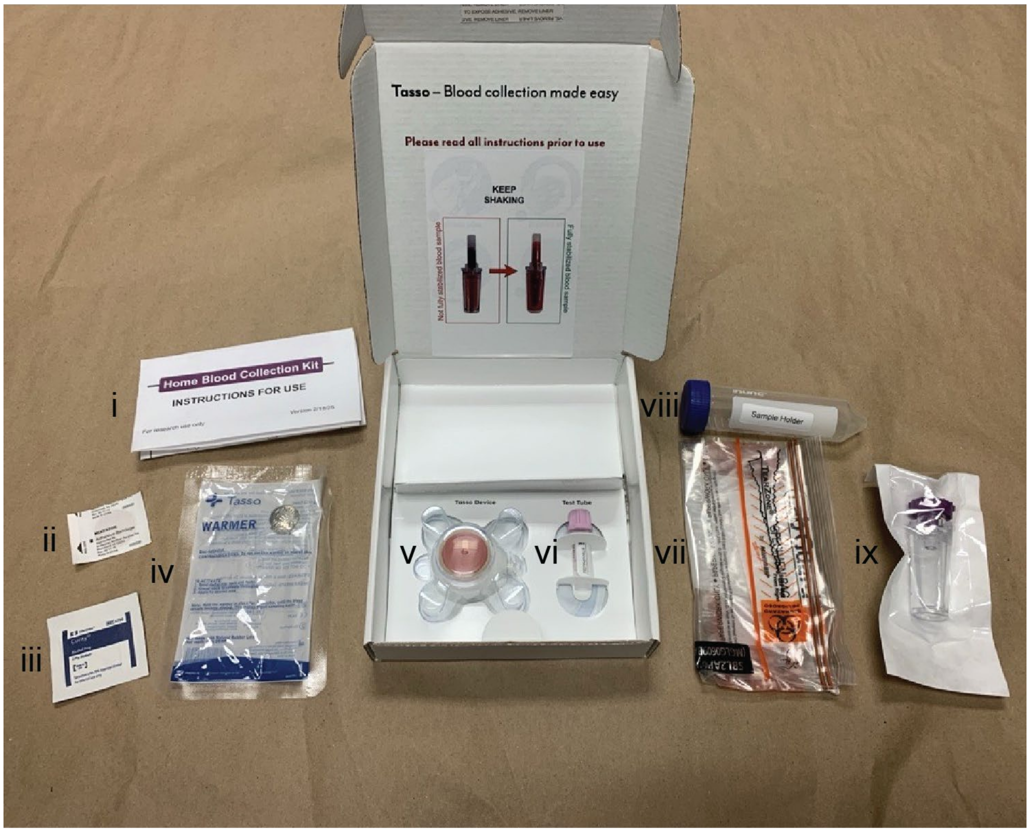


**Figure S6.** Components of the homeRNAmax kit. Kit components include i) instructions for use, ii) sterile bandage, iii) alcohol wipe, iv) warmer for warming the arm prior to application of the Tasso+ device, v) Tasso+ device, vi) BD Microtainer® blood tube, vii) sample return bag, viii) sample holder, and ix) stabilizer tube containing RNA*later*.

**Table S1. Components of the homeRNAmax kit.**

| Kit component | Manufacturer(s) | Quantity |
|---|---|---|
| Sterile Tasso+ blood collection device | Tasso, Inc. | 1 |
| RNA stabilizer tube | Our lab | 1 |
| BD Microtainer® blood tube (K2EDTA coated) | BD | 1 |
| Instant heat pack (product number ACC0014) | Tasso, Inc. | 1 |
| Sterile alcohol wipe | Covidien | 1 |
| Sterile bandage | McKesson | 1 |
| Specimen transport bag with absorbent pad | Path-Tec | 1 |
| 50 mL conical tube | BD | 1 |
| Instructions for use | Our lab (see page S6-S7 of SI) | 1 |
| Blood-stabilizer mixing instruction sticker | Our lab (see page S8 of SI) | 1 |

**Table S2.** homeRNAmax does not leak upon inversion at the volumes tested.*

| Volume of fluid in tube (mL) | Replicate | Height of fluid before inversion (mm) | Height of fluid above nozzle after inversion (mm) | Result |
|---|---|---|---|---|
| 2.00 | 1 | 11.0 | 1.99 | No leak |
| 2.00 | 2 | 13.2 | 4.11 | No leak |
| 2.00 | 3 | 12.9 | 3.09 | No leak |
| 2.00 | 4 | 12.9 | 3.27 | No leak |
| 4.00 | 1 | 22.6 | 11.5 | No leak |
| 4.00 | 2 | 24.9 | 13.7 | No leak |
| 4.00 | 3 | 24.1 | 13.3 | No leak |
| 4.00 | 4 | 23.4 | 13.1 | No leak |
| 6.00 | 1 | 32.8 | 22.5 | No leak |
| 6.00 | 2 | 34.0 | 24.3 | No leak |
| 6.00 | 3 | 33.7 | 24.3 | No leak |
| 6.00 | 4 | 33.7 | 24.0 | No leak |
| Filled (~6.70 mL) | 1 | 35.8 | 26.1 | No leak |
| Filled (~6.70 mL) | 2 | 37.2 | 28.8 | No leak |
| Filled (~6.70 mL) | 3 | 36.3 | 27.0 | No leak |
| Filled (~6.70 mL) | 4 | 36.3 | 27.5 | No leak |

*Note: Experiment was performed in quadruplicate. Volume of fluid used in homeRNAmax for RNA stabilization applications is 3.60 mL.

**Table S3**. homeRNAmax RNA concentration, yield, and data on blood collection. The concentrations were obtained on the Qubit Flex and are multiplied by 50 µL (elution volume). For the final concentrations reported in the manuscript, the concentrations from elution 1 and elution 2 are added together.

| Sample ID | Elution number | Concentration (ng/µL) | Yield (ng) | Time on arm (min.) | Blood level | Approximate blood volume (µL) |
|---|---|---|---|---|---|---|
| IGMJ | 1 | 32.4 | 1620 | >5 | Level E | 750 µL |
| | 2 | 4.11 | 205.5 | | | |
| AXUI | 1 | 16.1 | 805 | 3-5 | Level D | 500 µL |
| | 2 | 2.88 | 144 | | | |
| FJZM | 1 | 8.25 | 412.5 | 3-5 | Level A | 125 µL |
| | 2 | 1.3 | 65 | | | |
| EYWH | 1 | 33.7 | 1685 | 3-5 | Level D | 500 µL |
| | 2 | 3.48 | 174 | | | |
| CUBY | 1 | 29.3 | 1465 | 3-5 | Level E | 750 µL |
| | 2 | 3.87 | 193.5 | | | |
| BWNG | 1 | 42.3 | 2115 | 3-5 | Level F | 1000 µL |
| | 2 | 5.84 | 292 | | | |
| MPWJ | 1 | 87.8 | 4390 | 3-5 | Level F | 1000 µL |
| | 2 | 9.45 | 472.5 | | | |
| QLGG | 1 | 9.85 | 492.5 | 3-5 | Level F | 1000 µL |
| | 2 | 3.47 | 173.5 | | | |
| VIVZ | 1 | 2.15 | 107.5 | 3-5 | Level E | 750 µL |
| | 2 | 0.866 | 43.3 | | | |
| PLFF | 1 | 36.7 | 1835 | 3-5 | Level A | 125 µL |

| | | | | | | |
|---|---|---|---|---|---|---|
| PLFF | 2 | 6.21 | 310.5 | | | |
| SQFE | 1 | 12 | 600 | 3-5 | Level A | 125 µL |
| | 2 | 2.49 | 124.5 | | | |
| MQRF | 1 | 52.5 | 2625 | 3-5 | Level E | 750 µL |
| | 2 | 9.69 | 484.5 | | | |
| VHJF | 1 | 53.5 | 2675 | >5 | Level F | 1000 µL |
| | 2 | 10 | 500 | | | |
| POXS | 1 | 17.9 | 895 | 3-5 | Level C | 375 µL |
| | 2 | 4.62 | 231 | | | |
| VVXN | 1 | 26.1 | 1305 | 3-5 | Level F | 1000 µL |
| | 2 | 5.09 | 254.5 | | | |
| ZBVY | 1 | 78.8 | 3940 | <3 | Level F | 1000 µL |
| | 2 | 9.67 | 483.5 | | | |
| VYFO | 1 | 36 | 1800 | 3-5 | Level F | 1000 µL |
| | 2 | 9.64 | 482 | | | |
| ZIBT | 1 | 46.2 | 2310 | 3-5 | Level F | 1000 µL |
| | 2 | 9.97 | 498.5 | | | |
| PLFF-2 | 1 | 6.11 | 305.5 | >5 | Level E | 750 µL |
| | 2 | 4.34 | 217 | | | |

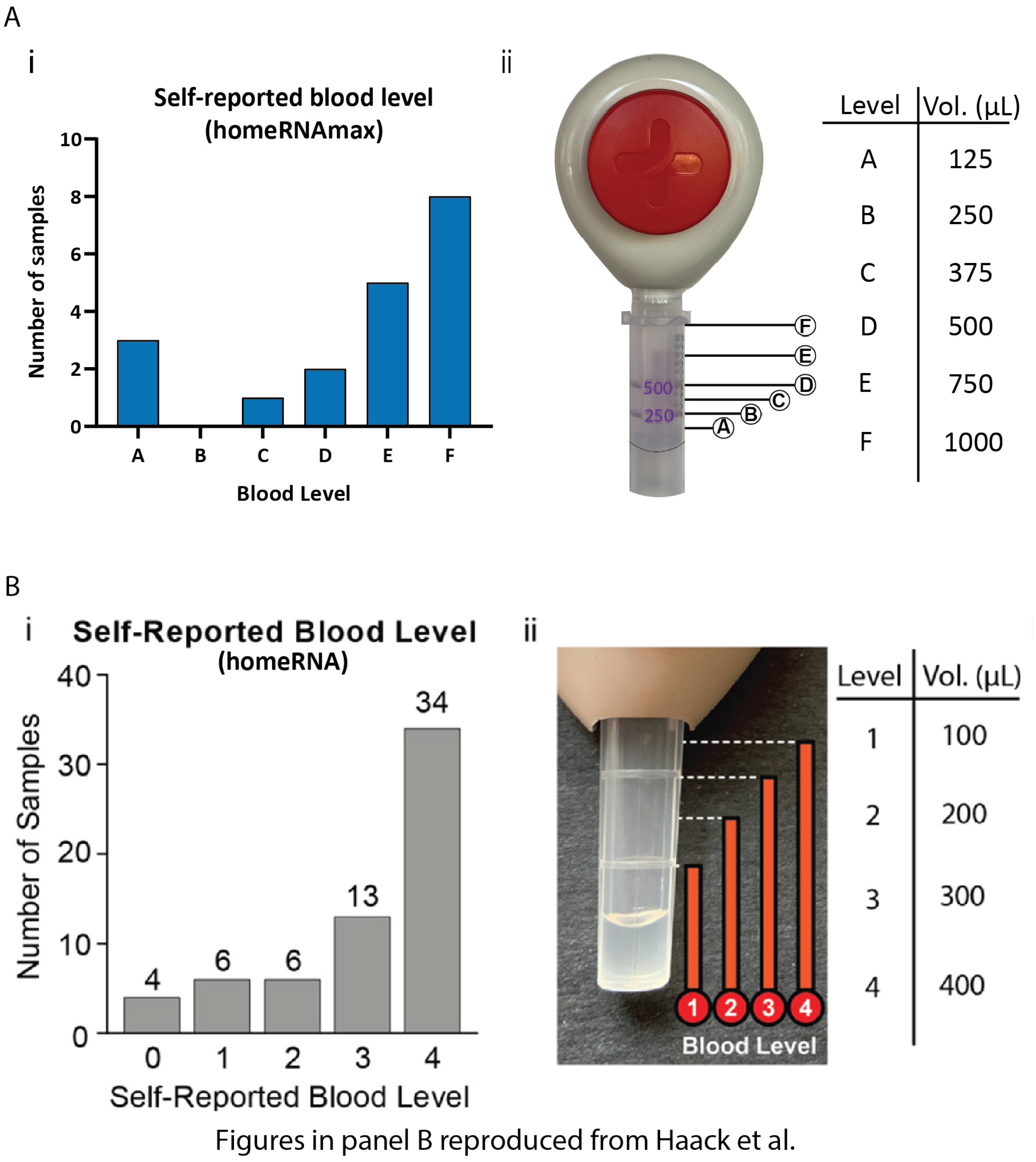


**Figure S7.** Comparison of homeRNAmax blood level (A, top) and original homeRNA[2] blood level (B, bottom) from participant-reported data following sampling. In the homeRNAmax, level B represents a volume of around 250 µL and level D represents a volume of 500 µL. The Images with annotated levels are included in the participant surveys as guidance for reporting collected blood level.

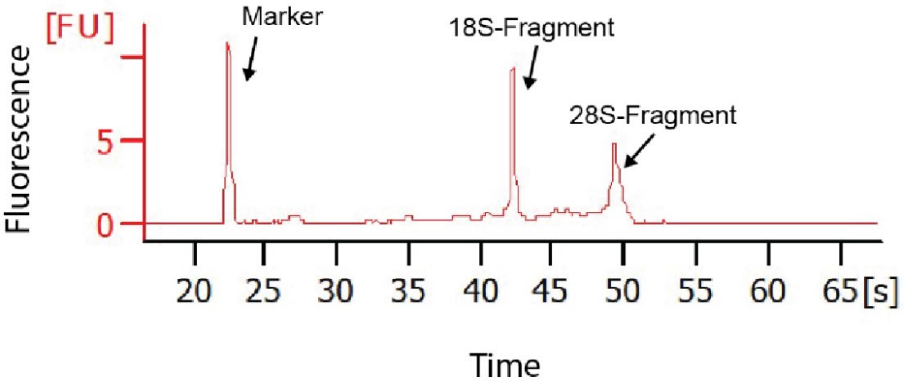


**Figure S8**. General Information on Bioanalyzer Profile. Electrophoretogram obtained from Sample B1 annotated with the marker and two other major peaks that help determine RIN including the 18-S fragment peak, and the 28-S fragment peak. More information on interpreting bioanalyzer data, including examples of electrophoretograms obtained from samples with various RIN values can be found in Schroeder 2006.[1] Figure and caption reproduced from Haack, A. J.; Lim, F. Y, et. al.[2]

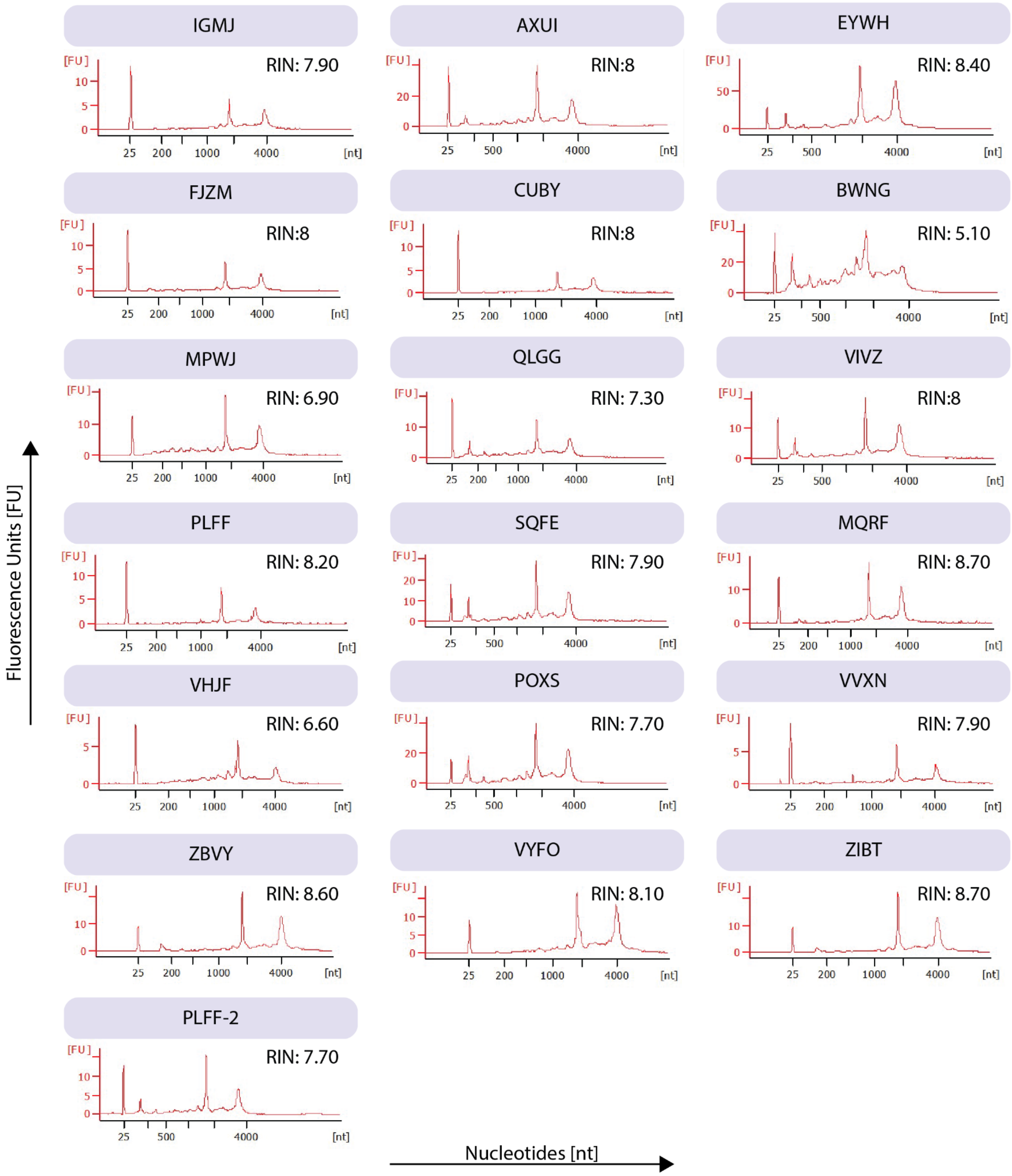


**Figure S9**. Agilent 2100 Bioanalyzer electropherograms for isolated RNA from homeRNAmax-stabilized samples. The samples were analyzed with Nano or Pico kits, depending on concentration. IGMJ, FJZM, CUBY, MPWJ, PLFF, MQRF, VHJF, VVXN, ZBVY, VYFO, and ZIBT were analyzed using the Nano kit, and AXUI, EYWH, BWNG, QLGG, VIVZ, SQFE, POXS, and PLFF-2 were analyzed using the Pico kit. For samples that were analyzed multiple times, the lowest of the two, or middle value of three is used and reported here and in the manuscript.

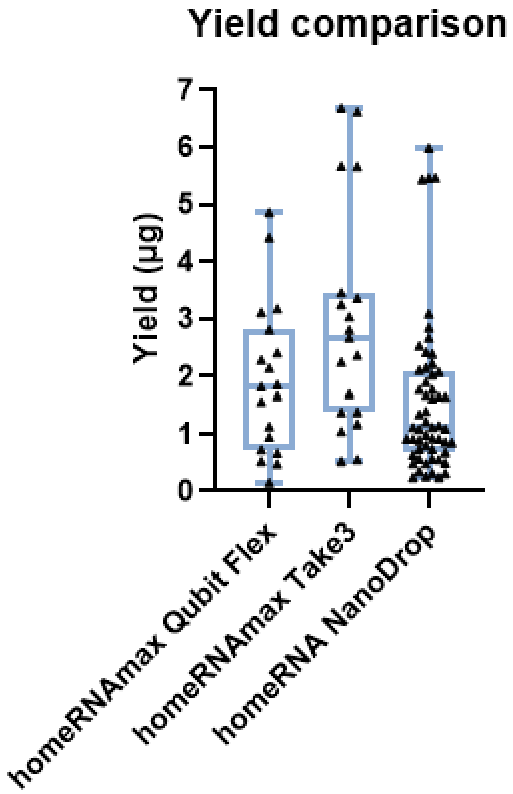


**Figure S10**. Comparison of RNA yield from present study (Qubit Flex, Cytation 5 Take3) and original homeRNA study[2] (NanoDrop). As expected, yields obtained from the Qubit Flex are slightly lower than those obtained from the Take 3. While an ideal comparison between the original homeRNA study and the present study is difficult because of the discrepancy in analytical instruments used, the Take3 and NanoDrop both use spectrophotometric methods (absorbance at 260 nm) to estimate RNA yield. When comparing these, it is clear that the distribution of the yields is similar between the homeRNA and homeRNAmax.

#### Appendix 1: Theoretical characterization of fluid retention in homeRNAmax stabilizer tube

In this study, we are analyzing the retention of a liquid in a homeRNAmax stabilizer tube tilted upside down (Fig. 1).

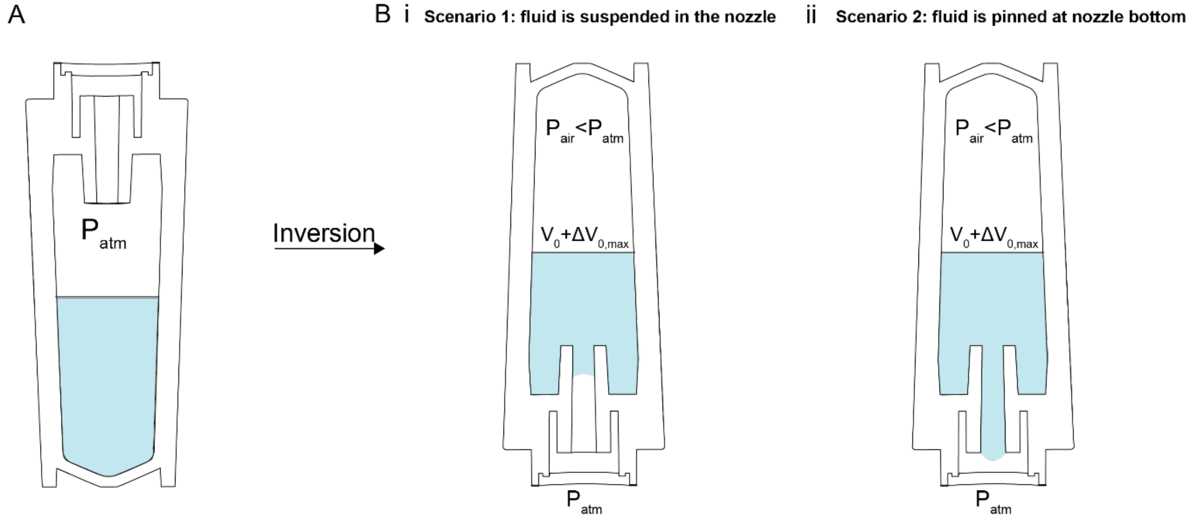


**Figure S11**. Schematic of stabilizer tube inversion. A) The tube before rotation. B) Schematic of an inverted stabilizer tube at equilibrium when i) fluid stops in the nozzle at equilibrium and ii) fluid is pinned to the nozzle opening at equilibrium. $P_{air}$ is the pressure of the air in the stabilizer tube, $P_{hyd}$ is the hydrostatic pressure of the fluid, $P_{cap}$ is the capillary pressure at the interface between the fluid and the atmosphere, and $P_{pin}$ is the pinning pressure.

#### Model

#### Scenario 1 (Fig. 1Bi): fluid is suspended in the nozzle

Inside the tube, we have the hydrostatic pressure ($P_{hyd}$), the pressure of the trapped air ($P_{air}$), and the capillary pressure in the nozzle ($P_{cap}$). The capillary pressure of the upper surface of the liquid in the tube is negligible since the tube diameter is larger than the capillary length. Outside the tube, the pressure is the atmospheric pressure ($P_{atm}$). The total pressure acting upon the meniscus suspended in the nozzle can be expressed by the following equation

$$P_{balance} = P_{hyd} + P_{cap} + P_{air} - P_{atm} \quad \text{(S1)}$$

The liquid is at equilibrium (immobile) when the pressure balance is zero

$$P_{hyd} + P_{cap} + P_{air} - P_{atm} = 0 \,. \quad \text{(S2)}$$

The hydrostatic pressure $P_{hyd}$ can be calculated using the following expression

$$P_{hyd} = \rho g\, H \quad \text{(S3)}$$

Where $\rho$ is the density of the fluid, $g$ is the acceleration due to gravity, and $H$ is the height of the fluid. In this case the pressure at the liquid air interface at the nozzle opening is a capillary pressure in the nozzle $P_{cap}$

$$P_{cap} = 2\gamma \cos\theta / r_{noz}. \quad \text{(S4)}$$

Where $\gamma$ is the surface tension of the fluid, $\theta$ is the contact angle of the fluid on the nozzle material, and $r_{noz}$ is the radius of the nozzle. Note that $P_{cap}$ is the same sign as $P_{hyd}$, meaning the capillary pressure does not prevent leakage in the tube.

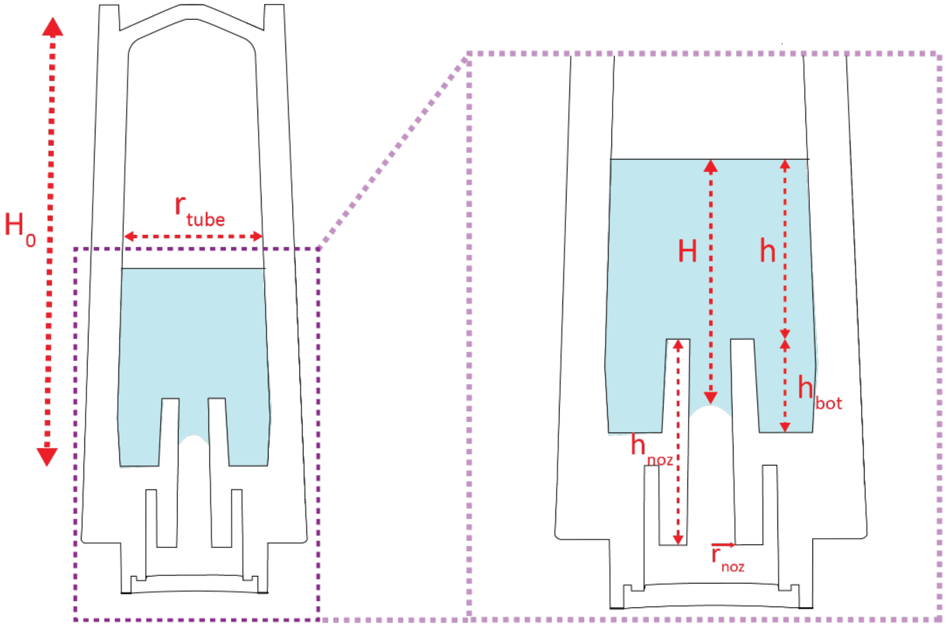


**Figure S12.** Schematic of stabilizer tube dimensions used in the theoretical model for Scenario 1.

By rearranging equation (S2), the pressure difference associated with the suspension of the fluid within the nozzle that is required to prevent leakage can be calculated.

$$P_{air} - P_{atm} = \Delta P_{air} = -(P_{hyd} + P_{cap}). \quad \text{(S5)}$$

It is worth noting that the precise pressure of the trapped air is difficult to determine. It depends on the way the tube has been tilted and on the initial volume in the tube in the upright position. It is likely that the liquid flows unto the extremity of the nozzle during the tilt. The suspension of the fluid in the nozzle decreases the pressure of the trapped air, initially taken as $P_{atm}$. If we apply the Ideal Gas Law we find:

$$P_{air} \sim P_{atm} \frac{V_0}{V_{air}} \quad \text{(S6)}$$

Where $V_0$ is the initial volume of air (before tilting) and $V_{air}$ is the volume of air after tilting. We can solve for the depression associated with the tilt by rearranging (S6):

$$\Delta P_{air} \sim P_{atm}\left(\frac{V_0}{V_{air}} - 1\right) \tag{S7}$$

Which can then be rearranged in terms of the change of volume:

$$\Delta P_{air} \sim -P_{atm}\left(\frac{\Delta V}{V_{air}}\right) \tag{S8}$$

The small amount of fluid that fills the nozzle would be sufficient to prevent leakage. Using water as an example, we can use equations (S3), (S4), and (S7) to approximate the order of magnitude of $\frac{\Delta V}{V_{air}}$ required to prevent leakage. If a stabilizer tube were filled with a height of water on the order of $10 - 10^2$ mm, the required pressure difference would be on the order of $10^2 - 10^3$ Pa. Since $P_{atm}$ is on the order of $10^5$ Pa, the difference in volume of the air (i.e., the volume of fluid that fills the nozzle) would only have to be on the order of $10^{-3}$.

#### Scenario 2 (fig. 1Bii): fluid is pinned at nozzle bottom

In this scenario, rather than fluid suspending in the nozzle, a droplet forms at the nozzle tip, and whether the drop leaks or stays pinned is governed by the sign of the function

$$P_{balance} = P_{hyd} - P_{pin} + P_{air} - P_{atm} \quad . \tag{S9}$$

If $P_{balance} > 0$ leakage occurs. We shall analyze what occurs when $P_{balance} < 0$. The expression for $P_{hyd}$ is the same as (S3), and, because the pendant droplet has a semi-spherical shape, $P_{pin}$ can be calculated using the following expression

$$P_{pin} = -2\gamma / r_{noz} \ . \tag{S10}$$

As for now, let us consider the value of the depression associated with the formation of the pendant drop. The volume of the pendant drop is

$$V_{drop} = 2/3\,\pi\, r_{noz}^3 \ . \tag{S11}$$

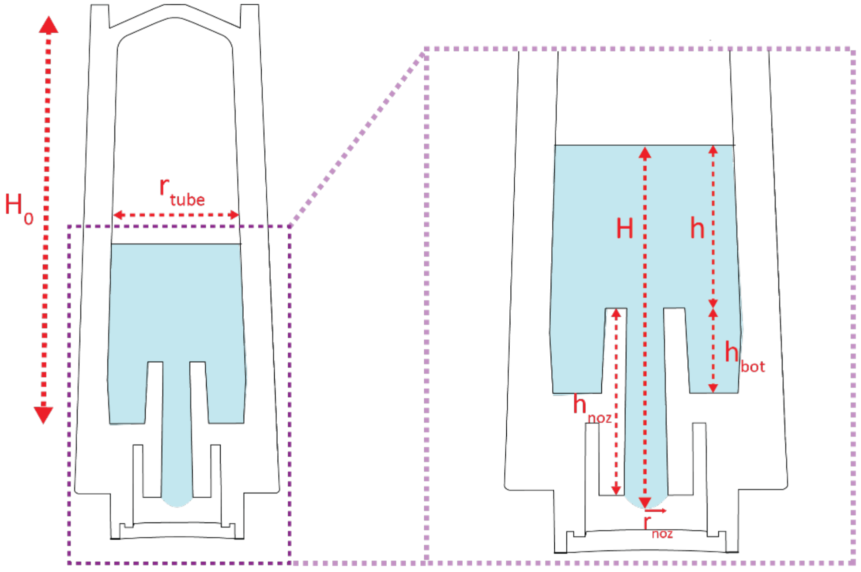

**Figure S13.** Schematic of stabilizer tube dimensions used in the theoretical model for Scenario 2. The variables are the same as in Scenario 1; they are repeated here for clarity.

So, the formation of the pendant drop decreases the pressure of the trapped air, initially taken as $P_{atm}$. If we apply the Ideal Gas Law

$$P_{air}(V_{air} + V_{drop}) = P_{atm}\, V_{air}\ . \tag{S12}$$

Then the term $P_{air} - P_{atm}$ becomes

$$P_{air} - P_{atm} = -P_{atm}\, \frac{V_{drop}}{V_{air}+V_{drop}}\ . \tag{S13}$$

So, the formation of the pendant drop creates a depression. If $V_{air}$ varies from nearly zero—if the tube is initially completely filled with liquid—to approximately the volume $V_{air} \approx H_0\, \pi r_{tube}^2$, the quantity $P_{air} - P_{atm}$ varies as shown in figure 4.

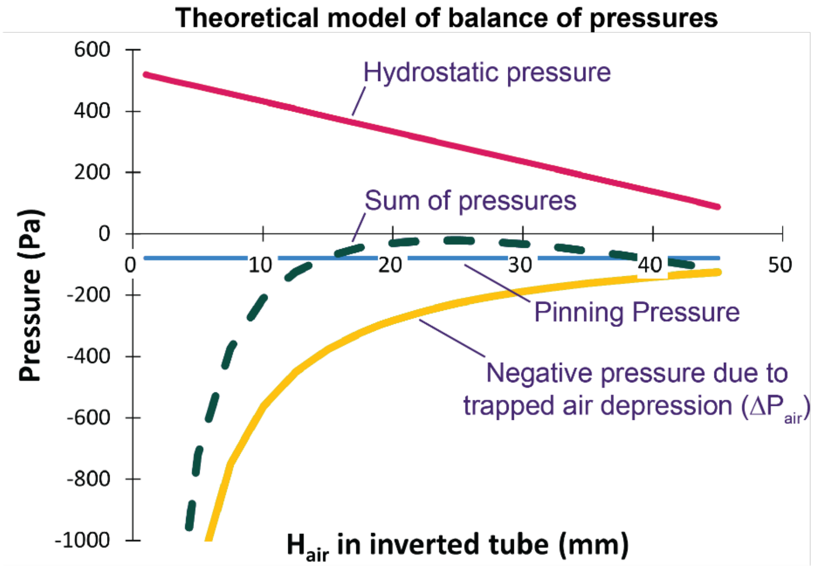


**Figure S14**. Theoretical prediction showing the sum of all pressures acting in the stabilizer tube (green) as a function of the height of the trapped air in the inverted stabilizer tube ($H_{air}$, mm), as well as the individual pressures (yellow, blue, and magenta).

When there is a small amount of liquid in the tube, the trapped air depression is small, and it is the pinning pressure that counterbalances the hydrostatic pressure. However, when the volume of liquid is large, the trapped air volume is small and the air depression becomes quite important. The pressure resultant $P_{balance}$ (green line in Fig. 4), has a maximum around $H_{air}$/15-25 mm.

#### Effect of a change of temperature on the equilibrium of the liquid in the tube

As noted at the end of *Theoretical model of the stabilizer tube* in the main text, some cases over timescales on the order of 10-20 minutes, additional meniscus movement has been observed after reaching equilibrium; we postulate that this movement is a result of effects related to temperature (e.g., evaporation, temperature changes over time). We have not observed any leakage as a result of these effects. Furthermore, in actual use cases, the tube is not left uncapped and inverted for longer than times on the order of seconds. The following section explores the effects of a temperature change.

Consider an inverted tube at equilibrium that undergoes a temperature change while inverted. The trapped air interface slightly moves upwards and the meniscus in the nozzle also moves upwards to reach another equilibrium state. Let us denote by the subscript 1 the first equilibrium state and subscript 2 the new equilibrium state.

The two equilibrium states are defined by

$$P_{air,1} + P_{hyd,1} + P_{cap} - P_{atm} = 0, \tag{S14}$$

And

$$P_{air,2} + P_{hyd,2} + P_{cap} - P_{atm} = 0, \tag{S15}$$

Then subtracting (S14) from (S15)

$$P_{air,2} - P_{air,1} = P_{hyd,1} - P_{hyd,2}. \tag{S16}$$

The first equilibrium hydrostatic pressure is determined by

$$P_{hyd,1} = \rho g(h + z), \tag{S17}$$

where *h* is the height of liquid above the upper end of the nozzle and *z* the height of liquid inside the nozzle. A decrease of pressure triggers the upwards motion of the meniscus of *Dz*, and a corresponding motion of the air/liquid surface *Dh*. Using the mass conservation equation

$$\Delta V = \pi r_{noz}^2 \Delta z = \pi r_{tub}^2 \Delta h. \tag{S18}$$

Then, the second equilibrium hydrostatic pressure is

$$P_{hyd,2} = P_{hyd,1} - \rho g \Delta z + \rho g \Delta h = P_{hyd,1} + \rho g \Delta h \left(1 - \frac{r_{tub}^2}{r_{noz}^2}\right). \tag{S19}$$

On the other hand, the trapped air volume decreases by

$$V_{air,2} = V_{air,1} - \pi r_{tub}^2 \Delta h\ . \tag{S20}$$

In terms of height

$$H_{air,2} = H_{air,1} - \Delta h\ . \tag{S21}$$

The new air pressure is then given by

$$\frac{P_{air,2}V_{air,2}}{T_2} = \frac{P_{air,1}V_{air,1}}{T_1}\,, \tag{S22}$$

so that

$$P_{air,2} = \frac{P_{air,1}V_{air,1}}{V_{air,2}}\frac{T_1}{T_2} = \frac{P_{air,1}V_{air,1}}{V_{air,1} - \pi r_{tub}^2 \Delta h}\frac{T_1}{T_2} = \frac{P_{air,1}H_{air,1}}{H_{air,1} - \Delta h}\frac{T_1}{T_2}\,, \tag{S23}$$

Substitution of (S19) and (S20) in (S16) yields

$$P_{air,2} - P_{air,1} = P_{air,1}\left(\frac{H_{air,1}}{H_{air,1} - \Delta h}\frac{T_1}{T_2} - 1\right) = P_{hyd,1} - P_{hyd,2} = \rho g \Delta h \left(\frac{r_{tub}^2}{r_{noz}^2} - 1\right). \tag{S24}$$

It is verified in (S24) that, if $T_2 = T_1$, then $\Delta h = 0.$ Because the first equilibrium state is known, $H_{air,1}$ and $P_{air,1}$ are known. We then obtain an equation in $\Delta h$. Let us cast (S24) under the form

$$P_{air,1}\left(\frac{1}{1 - \frac{\Delta h}{H_{air,1}}}\frac{T_1}{T_2} - 1\right) = \rho g \frac{\Delta h}{H_{air,1}} H_{air,1}\left(\frac{r_{tub}^2}{r_{noz}^2} - 1\right), \tag{S25}$$

where $\varepsilon = \Delta h / H_{air,1}$ is now the unknown to determine. Then we obtain the quadratic equation

$$\frac{\rho g H_{air,1}}{P_{air,1}}\left(\frac{r_{tub}^2}{r_{noz}^2} - 1\right)\varepsilon^2 + \left[1 - \frac{\rho g H_{air,1}}{P_{air,1}}\left(\frac{r_{tub}^2}{r_{noz}^2} - 1\right)\right]\varepsilon + \frac{T_2}{T_1} - 1 = 0\,. \tag{S26}$$

Again, if $T_2 = T_1$, $\varepsilon = 0$ is solution, and $\Delta h = 0$. Note that $\frac{\rho g H_{air,1}}{P_{air,1}}$ is non-dimensional so that (13) is non-dimensional. An approximate solution can be obtained if the term in $\varepsilon^2$ is negligible, and

$$\varepsilon = \frac{\Delta h}{H_{air,1}} \approx \frac{\frac{T_2}{T_1} - 1}{\left[1 - \frac{\rho g H_{air,1}}{P_{air,1}}\left(\frac{r_{tub}^2}{r_{noz}^2} - 1\right)\right]}\,. \tag{S27}$$

Let us consider $T_2 = 20\,C = 293\,K$, $T_1 = 28\,C = 301\,K$, $T_2/T_1 \approx 0.97$, $r_{tub}/r_{noz} = 3$, $H_{air,1} = 20\,mm$, $P_{air,1} - P_{atm} = 300\,Pa$, so that $P_{air,1} = 9700\,Pa$. The term $\frac{\rho g H_{air,1}}{P_{air,1}}$ is equal to $2 \times 10^{-3}$ approximately. Then $\varepsilon = \frac{\Delta h}{H_{air,1}} \approx \frac{\frac{T_2}{T_1} - 1}{\left[1 - \frac{\rho g H_{air,1}}{P_{air,1}}\left(\frac{r_{tub}^2}{r_{noz}^2} - 1\right)\right]} \approx 0.03$. Hence $\Delta h \approx 0.6\,mm$ and $\Delta z \approx 4.8\,mm$
