## Supplementary Information 2 for "To the homeRNAmax: Developing an Improved Blood Self-Collection and Stabilization Platform for Remote Transcriptomic Studies": homeRNAmax-Kit-Survey.pdf

### homeRNA+ Kit Survey

Please complete the survey below.

Thank you!

---

What is the kit code?

__________________________________

---

Did you experience any issues with the Tasso+ blood collection device?

- ◯ Yes
- ◯ No

---

Please describe any issues you had with the Tasso+ blood collection device:

__________________________________________

---

Did you experience any pain when using the homeRNA+ kit today?

- ◯ No pain
- ◯ Mild pain
- ◯ Moderate pain
- ◯ Severe pain
- ◯ Very severe pain

---

Were you able to collect any blood?

- ◯ Yes
- ◯ No

---

If you have any comments or suggestions for the instructions, please use this space:

__________________________________________

---

Approximately how long (in minutes) did it take you to use the homeRNA+ blood kit today?

- ◯ Less than 5 minutes
- ◯ 5-10 minutes
- ◯ 11-15 minutes
- ◯ 16-20 minutes
- ◯ More than 20 minutes

---

While using the homeRNA+ blood kit today, approximately how long (in minutes) did you leave the Tasso+ blood collection device on your arm?

- ◯ Less than 1 minute
- ◯ Less than 3 minutes
- ◯ 3-5 minutes
- ◯ More than 5 minutes
- ◯ I am unsure

---

Based on the image below, how much blood did you roughly collect today? Choose the closest Level.

- ◯ Level A
- ◯ Level B
- ◯ Level C
- ◯ Level D
- ◯ Level E
- ◯ Level F

---

Did you experience any issues with the stabilizer tube or mixing your sample?

- ◯ Yes
- ◯ No

---

Please describe any issues you had with the stabilizer tube or mixing your sample:

__________________________________________

**How easy was it to use the components of the kit listed below?**

| | Very difficult | Somewhat difficult | Neither difficult nor easy | Somewhat easy | Very easy |
|---|---|---|---|---|---|
| Kit Instructions | ○ | ○ | ○ | ○ | ○ |
| The Tasso+ device | ○ | ○ | ○ | ○ | ○ |
| Blood Stabilization Tube | ○ | ○ | ○ | ○ | ○ |

Do you have any other thoughts or comments regarding your experience with this kit? (optional)

__________________________________________
