## Supplementary Information 2 for "To the homeRNAmax: Developing an Improved Blood Self-Collection and Stabilization Platform for Remote Transcriptomic Studies": homeRNAmax_emailed study instructions.pdf

Hi [prefname],

Now that you have received your kit, you are ready to collect your sample and take the survey!

Please complete the following steps within 1 - 2 days after receiving this email:

**1. Collect the Sample**

- Please watch the provided video before you begin sampling
  - Video is linked to QR code in physical instructions, and can be accessed through this link
- Follow printed instructions to collect your sample

**2. Prepare Sample for Shipping**

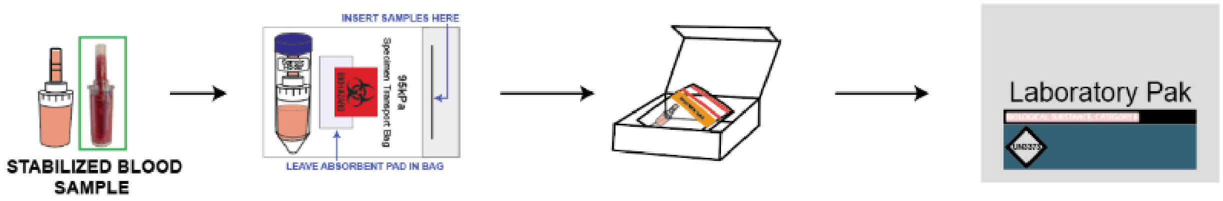


- Once the sample is secured in the kit box, place it inside the pre-labeled UPS LabPak
  - Seal LabPak by removing the adhesive strip and pressing the flaps together
- Keep sample indoors overnight at room tempearture
- Place LabPak at the designated pickup location the following morning by 9:00am
  - If the next day is Sunday, please leave package inside until Monday morning

Note: there is nothing you need to do to schedule the pickup, we automatically schedule them after you complete the survey below. You will receive an additional email letting you know what day the pickup is scheduled for.

**3. Complete Survey**

Finally, please complete this survey after you are done with the sample collection:

[survey-link]

If the link above does not work, try copying the link below into your web browser:
[survey-url]

This link is unique to you and should not be forwarded to others.

Thank you again for your participation and please let us know if you have questions or need assistance!

All the best,

BCME Study Team

University of Washington
