## Supplementary figures and images for "To the homeRNAmax: Developing an Improved Blood Self-Collection and Stabilization Platform for Remote Transcriptomic Studies"

### homeRNAmax_inside_sticker.pdf

**KEEP SHAKING**

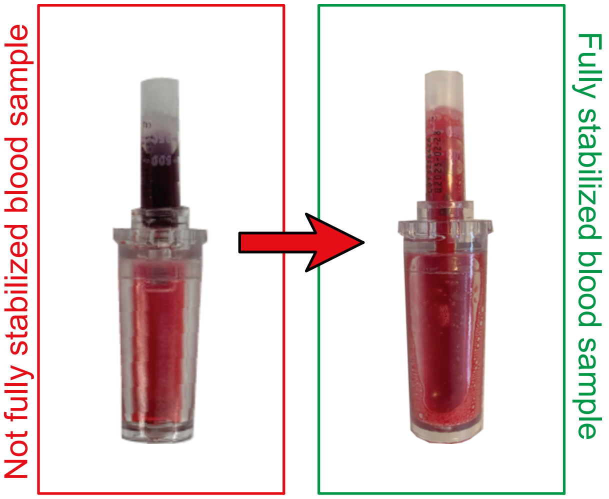


**KEEP SHAKING**

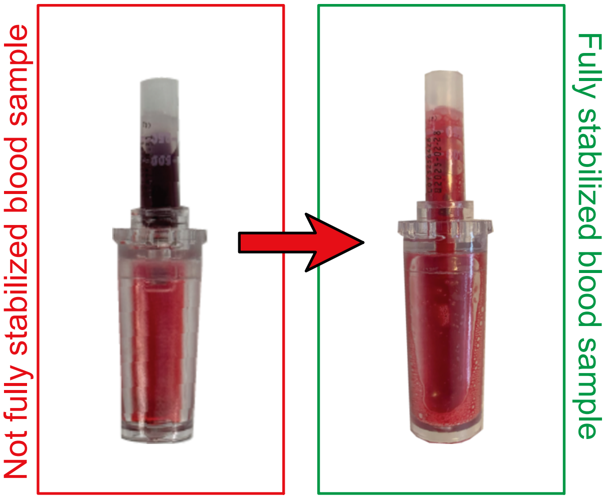


**KEEP SHAKING**

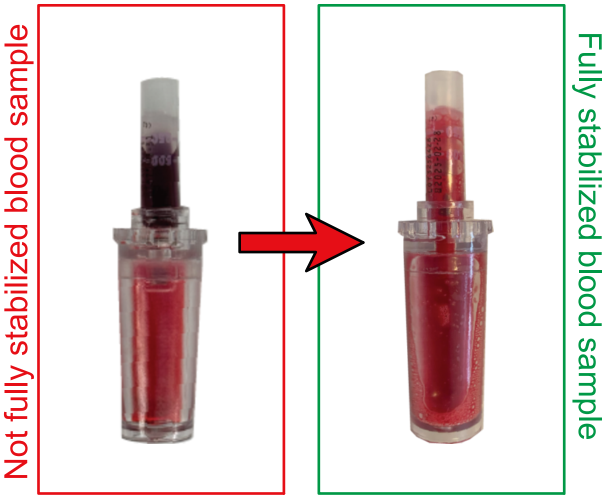


**KEEP SHAKING**

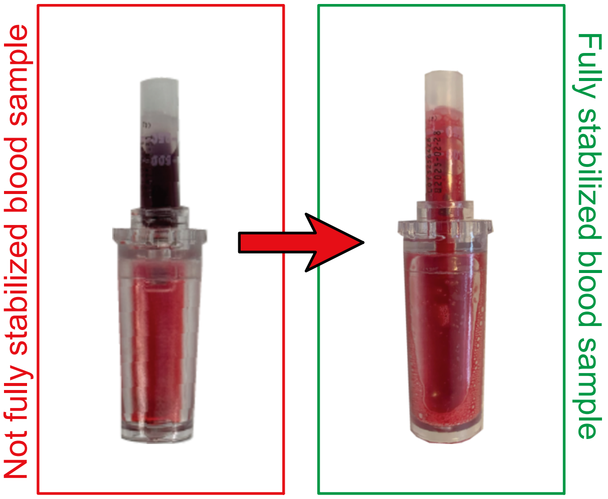
